# Whole-genome sequence of *Potamopyrgus antipodarum*—a model system for the maintenance of sexual reproduction—reveals a recent whole-genome duplication

**DOI:** 10.1101/2025.03.16.643514

**Authors:** Joseph Jalinsky, Kyle McElroy, Joel Sharbrough, Laura Bankers, Peter Fields, Chelsea Higgins, Cynthia Toll, Jeffrey L. Boore, John M. Logsdon, Maurine Neiman

## Abstract

Key unanswered questions in biology center on the causes, consequences, and maintenance of sexual reproduction (“sex”). Genome-driven processes are central to the evolutionary and genetic mechanisms inherent to sex, making genomic resources a fundamental part of answering these questions. We present the first genome assembly for a species that is uniquely well-suited for the study of (a)sex in nature, *Potamopyrgus antipodarum*. This New Zealand snail is unusual in featuring multiple separate transitions from obligately sexual to obligately asexual reproduction, leading to the coexistence of phenotypically similar sexual and asexual forms, a feature that is required to directly study the maintenance of sex. These separately derived asexual lineages constitute separate evolutionary experiments, providing a powerful means of characterizing how the absence of sex affects genome evolution. Our genome assembly provides critical steps towards understanding causes and consequences of sex in this system and important resources for the rapidly growing *P. antipodarum* and molluscan genomics research community. In characterizing this genome, we uncovered unexpected evidence for a recent whole-genome duplication (WGD) in *P. antipodarum*. This discovery sets the stage for using *P*. *antipodarum* to evaluate processes of rediploidization following WGD and assess whether WGD might drive transitions to asexuality.

## Introduction

The overwhelming predominance of sexual reproduction in eukaryotes has fascinated biologists for at least 150 years, when Charles Darwin (1862) pointed out that any explanation for why so many organisms use sex instead of parthenogenesis was “hidden in darkness” [1]. While we have since learned much about *how* organisms sexually reproduce (reviewed in [2,3]), the *why* remains a remarkably persistent evolutionary problem [4–8].

In principle, it should be simple to identify the drivers of the maintenance of sexual reproduction by comparing factors thought to be related to advantages of sex (*e.g.*, frequency of intense parasitism, extent of resource limitation) across individuals, populations, or lineages that differ in reproductive mode. In practice, these comparisons are very challenging, in large part because polymorphism for sexual and asexual reproduction in otherwise similar and coexisting individuals is rare in natural populations [8,9]. The absence of such polymorphism means that the maintenance of sex is not at stake and implies that studies of systems that do not feature conspecific, sympatric, and otherwise similar sexual and asexual individuals are only indirectly applicable to the sex question [9].

### *Potamopyrgus antipodarum* is a powerful model for the study of (a)sex in nature

Populations of the ancestrally sexual and dioecious Aotearoa New Zealand (hereafter “New Zealand”) freshwater snail *Potamopyrgus antipodarum* are characterized by coexistence of phenotypically indistinguishable obligate sexuals and obligate asexuals [10], enabling direct comparisons between sexual and asexual individuals and lineages and sexually and asexually propagated genomes. The relative frequency of sexual *vs*. asexual *P. antipodarum* is temporally stable within lakes but varies widely across lakes [11], enabling the population comparisons required to identify environmental factors contributing to intraspecific maintenance of reproductive mode polymorphism (*e.g.*, [10,12,13]).

Asexual lineages of *P. antipodarum* have been derived on many separate occasions from sexual *P. antipodarum* [14–16], allowing us to treat each distinct lineage as a replicated test of asexuality. The *P. antipodarum* system is especially unusual in that asexual lineage origin does not appear to be associated with hybridization [9,14,17]. This phenomenon suggests an intrinsic predisposition towards transitions to asexual reproduction (reviewed in [18]).

*Potamopyrgus antipodarum* is therefore especially well-suited to apply to major outstanding questions in biology regarding the genetic and genomic factors driving transitions to obligate asexual reproduction, the role of genetic and genomic mechanisms in the maintenance of sexual reproduction, and the genomic consequences of asexual reproduction. *P*. *antipodarum* is also a textbook example for host-parasite coevolution [10,19,20] and is used as a model for ecotoxicology [21,22], mitochondrial-nuclear coevolution [23–27], invasion biology [28–31], and polyploidy [16,32,33], indicating that the resources described here will be of wide use.

Here, we report the first draft genome assembly for *P. antipodarum* (Figure 1). We also uncover unexpected and striking evidence for a recent and still-resolving whole-genome duplication (WGD) event that provides a unique opportunity to study the ongoing process of rediploidization and may help explain the high frequency of transitions to obligate asexual reproduction in this species. The availability of a new high-quality molluscan genome assembly will also be of broad interest and utility to the wider community of molluscan genomics researchers [34].

**Figure 1.**
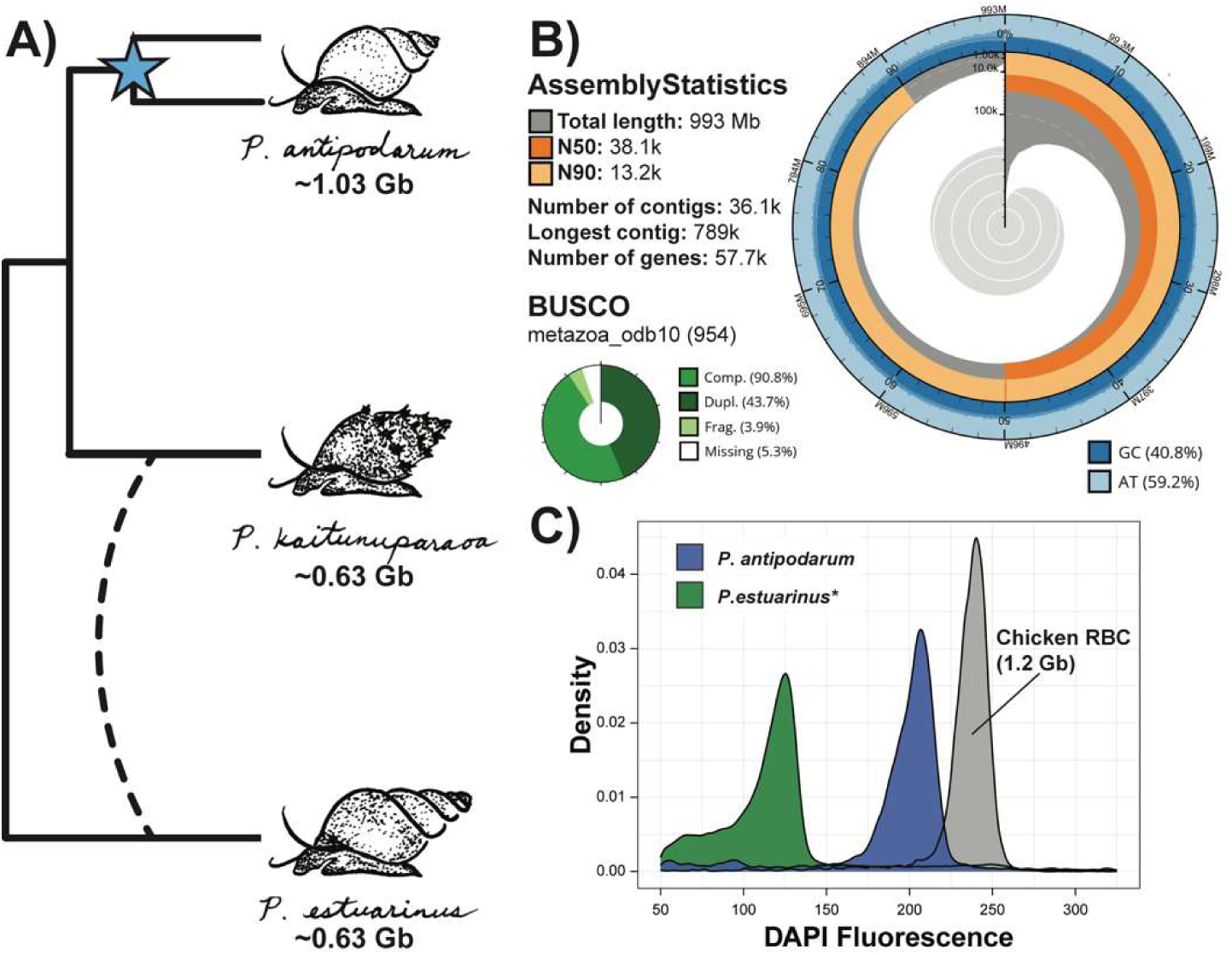
The genome assembly of *Potamopyrgus antipodarum*. A) Cladogram depicting species relationships among *Potamopyrgus* species. Below each species name is the approximate haploid genome size estimated from flow cytometry. The two branches leading to *P. antipodarum* represent the apparent whole-genome duplication event (blue star) within the species. The dashed line represents putative introgression inferred between *P. kaitunuparaoa* and *P. estuarinus*. See [50] and Supplementary Figure S6 for additional details on this introgression, which is not the main focus of this paper. B) Snail plot describing assembly statistics for the “uncollapsed” *P. antipodarum* contigs. The snail plot in the center is divided into 1,000 size-ordered bins, with each bin representing 0.1% of the assembly. Sequence length distribution is shown in dark gray, with the radius scaled to the longest sequence (shown in red). The N50 sequence length (orange) and N90 sequence length (pale orange) are represented by the arcs in the center of the plot. The cumulative sequence count is represented by the pale gray spiral, with white scale lines showing orders of magnitude. GC and AT percentages are reflected by the dark blue and pale blue areas, respectively, around the outside of the plot. BUSCO marker summary statistics are shown in the bottom left. C) Estimated genome size of *P. antipodarum* (blue) and *P. estuarinus* (green) from DAPI-stained nuclei, compared to chicken red blood cells (grey). Asterisk refers to a similar genome size for *P. kaitunuparaoa* separately estimated using propidium iodide (data not shown).

## Methods

To assemble a representative genome sequence from *P. antipodarum*, we generated genomic data from an inbred sexual diploid lineage (“Alex Yellow”) founded by a sexual female collected from Lake Alexandrina in the 1990’s. We selected this lineage for the reference genome because it had been inbred for ∼20 years (∼20-30 generations; [35]) and was thus likely to have relatively high homozygosity, facilitating the genome assembly process. This lineage has also been used for other sequencing projects [24,26,36]. We used three additional Alex Yellow individuals for the generation of transcriptomic data: one brooding (reproductive) adult female, one non-brooding adult female, and one adult male. Detailed methods for assembly and annotation with MAKER and BUSCO [37] can be found in the Supplementary Information.

### Evaluating evidence for a recent WGD in *P. antipodarum*

*Genome size estimation with flow cytometry:* We determined approximate total nuclear genome size using flow cytometry, as described in [38]. Briefly, flash-frozen *P. antipodarum* and *P. estuarinus* head tissues were ground in a solution containing 0.2 M Tris-HCl (pH 7.5), 4 mM MgCl_2_, 1% TritonX-100, and 4 μg/mL DAPI. This solution was filtered through a 70-micron nylon sheet and assayed on a Beckman-Coulter Quanta SC MPL flow cytometer (Beckman Coulter, Brea, USA). Four *P. estuarinus* samples were assayed on January 27, 2015, and 40 *P. antipodarum* Alex Yellow samples were assayed on February 13, 2015. We used the FL1 channel to assess DAPI fluorescence (and thus the total DNA content) of cell nuclei under a UV lamp. At the beginning and end of each flow cytometry run, we calibrated the machine using 20 μL of chicken red blood cells (Lampire Biological Labs, Pipersville, PA) treated and filtered as for *Potamopyrgus* samples. Each sample was run until a count of at least 10,000 events was achieved.

*Allele sequencing depth at heterozygous sites:* As a preliminary test of WGD, we designed a mapping experiment to compare relative sequencing coverage of alleles at heterozygous sites. To do this, we mapped Illumina HiSeq reads collected from a single individual female snail to the collapsed assembly (see Supplementary Information) using bwa v0.7.17-r1188 [39] with default parameters. We called variants using GATK v4-4.6.1.0-0. As part of this procedure, we first removed PCR and optical duplicates using the MarkDuplicates function, then called variants, assuming diploidy, using the HaplotypeCaller function. We extracted allele sequencing depth from the DP4 field of the resulting VCF file for every heterozygous site called by GATK4, calculated the proportion of reads covering the minor allele as a fraction of the total depth over the site, and visualized the distribution of these data using the ggplot2 subpackage [40] in R v4.4.2 [41].

*Site-frequency spectrum analysis:* The distribution of allele frequencies (i.e., site-frequency spectrum; SFS) at biallelic sites can be used to infer genomic copy-number and, thus, estimate ploidy *in silico* [42–48]. Here, we used nQuire [49], a bioinformatic tool that estimates intragenomic SFS, to test for the possibility of a WGD event specific to the *P. antipodarum* lineage. As part of this analysis, we compared the nQuire output from *P. antipodarum* to that obtained from the assemblies of diploid congeners *Potamopyrgus estuarinus* and *Potamopyrgus kaitunuparaoa* [50]. Additional details about this analysis are available in the Supplementary Information.

*Gene-based tests for WGD:* The outcomes of the analyses described above, coupled with the challenges we experienced during *de novo* assembly of the *P. antipodarum* genome, led us to hypothesize that *P. antipodarum* had experienced a recent WGD. We evaluated this hypothesis through a series of gene-based analyses: 1) comparative sequence-based intragenomic homology (*i.e.*, global distribution of paralogs in *P. antipodarum vs. P. kaitunuparaoa*, and *P. estuarinus*), 2) phylogeny-based inference of paralog divergence and tree topology, and 3) synteny-based homology (*i.e.*, collinearity of duplicated genes in *P. antipodarum*). Details of these analyses are provided in the Supplementary Information.

## Results

We used four different genomic sequencing technologies and RNA sequencing to produce the first-ever high-quality reference genome assembly and annotation for *Potamopyrgus antipodarum*. This diversity of data was necessary because of the high complexity of the *P. antipodarum* genome, which, as we document below, is largely a consequence of a relatively recent WGD event (Figure 1A). Accordingly, we varied genome assembly parameters to produce two distinct genomic resources from these data, both of which will be of wide utility for the *Potamopyrgus* research community: (1) a more biologically realistic but substantially more fragmented set of ‘uncollapsed’ contigs that have not been scaffolded so as to best separate the duplicated regions of the genome, and (2) a computationally haploidized set of ‘collapsed’ scaffolds that provides large-scale contiguity and simplicity at the expense of structural reality.

### Assembly and annotation of the *P. antipodarum* genome

*Assembly and gene annotation:* Our uncollapsed *P. antipodarum* assembly had a contig N50 of 38.1 Kb and total length of 993 Mb (Figure 1B, Supplementary Figure S2A). The collapsed version of our *P. antipodarum* assembly had a scaffold N50 of 1.14 Mb and total length of 543 Mb (Supplementary Figure S2B). In total, we annotated 57,703 protein-coding genes in the uncollapsed contigs, 79.0% more genes than the *P. estuarinus* assembly (32,237) and 90.4% more genes than the (relatively incomplete) *P. kaitunuparaoa* assembly (30,310) [50]. In agreement with the uncollapsed assembly, DAPI-based flow cytometry indicated that *P. antipodarum* (∼2.06 pg) has a substantially larger (1.6 times, or 61%, larger) nuclear genome than *P. estuarinus* (∼1.26 pg) (Figure 1C, Supplementary Figure S1) and *P. kaitunuparaoa* [50,51]. Additional details about both assemblies and their annotation are provided in the Supplementary Information, and all materials have been made publicly available (see Data Availability statement).

*Meiosis gene inventory:* The presence of intact and complete meiosis-specific genes in the genome of an organism is consistent with the ability to engage in meiosis, and thus, sex [52]. We compared the meiosis gene repertoire of *P. antipodarum* to those of congeners *P. estuarinus* and *P. kaitunuparaoa* and found that the collapsed and uncollapsed *P*. *antipodarum* genome assembly contains the same complement of meiosis genes as in the other two species (38 of the 44 queried meiosis genes, Figure 2). The only exception, DMC1, was present in single copy in the uncollapsed contigs but was absent from the collapsed scaffolds. Every gene absence in the uncollapsed *P*. *antipodarum* assembly was mirrored in *P*. *estuarinus* and *P*. *kaitunuparaoa* [50], likely indicating true absence as opposed to an artifact of mis/non-assembly. Three meiosis genes (RECQ3, TIM2, and CycA) were absent from all three species, potentially indicating that these genes are not necessary for meiosis in these caenogastropods. Of the 38 meiosis genes found in the uncollapsed assembly, 23 (60%) were found to exist in multiple copies (single copy = 15/38; double copy = 15/38; triple copy = 8/38). The duplicated genes are non-identical to one another with one exception (PLK1) and have a nucleotide p-distance distribution ranging from 0 - 0.034 (x̅=0.011; SD = 0.009).

**Figure 2.**
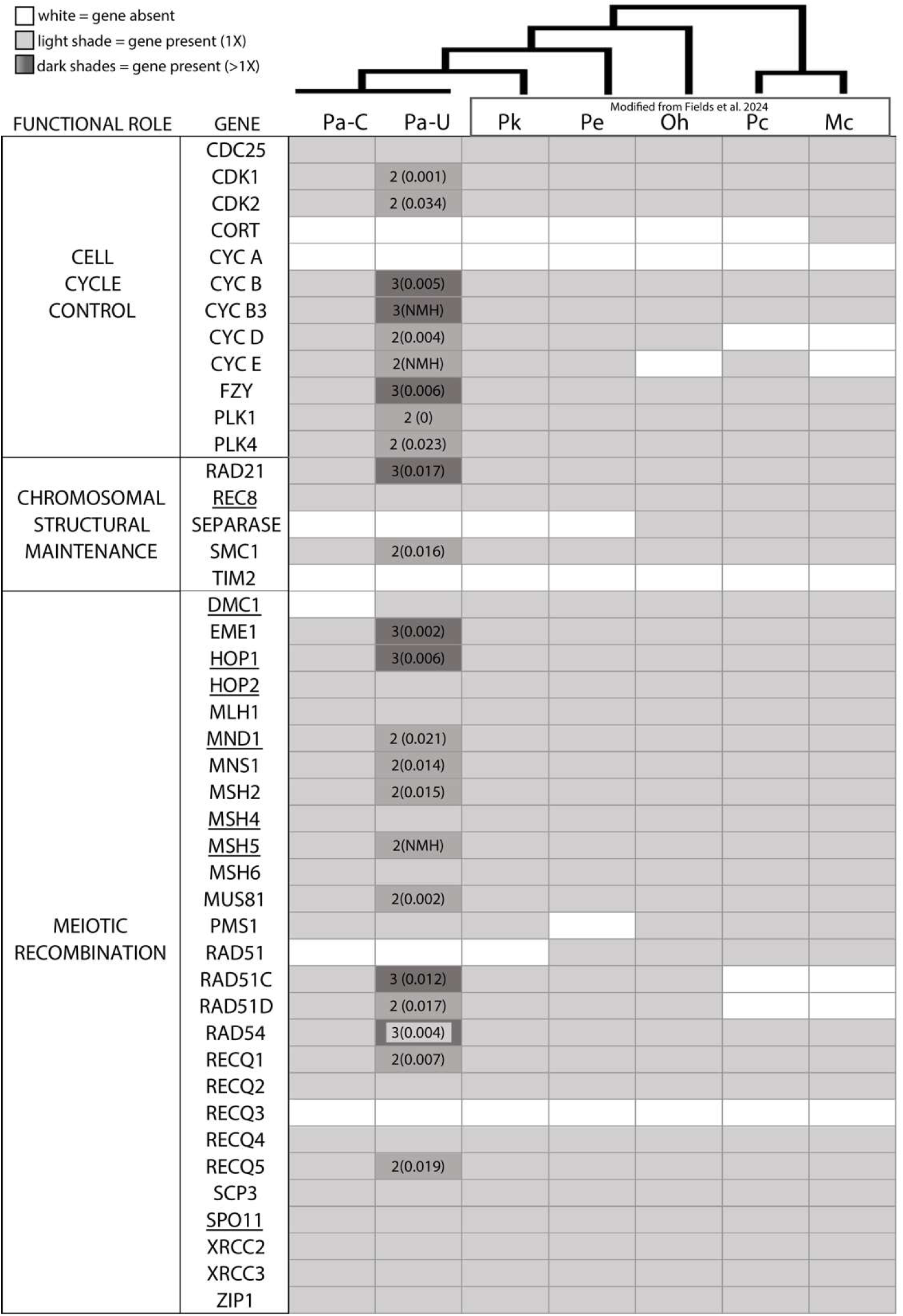
Gene inventory for 44 meiosis genes in six gastropod species. Meiosis gene presence and absence including data from this study and from [50]. *Pa* = *P. antipodarum* (the ‘C’ and ‘U’ represent the collapsed and uncollapsed genome assemblies, respectively); *Pk* = *P. kaitunuparaoa*; *Pe* = *P. estuarinus*; *Oh* = *Oncomelania hupensis*; *Pc* = *Pomacea canaliculata*; *Mc* = *Marisa cornuarietis*. For each cell, the number to the left of the parentheses indicates the number of copies present where the gene is at least 80% of the length of the query gene. The number inside the parentheses is the nucleotide p-distance of the mean nucleotide difference between the duplicated copies from the MAKER annotation. NMH = No MAKER hits (these genes were not recovered by the MAKER annotation but were discovered by BLAST and miniprot). A gene is present in each genome assembly if a box is shaded; if the shaded box is empty the gene is present in single copy. A gene is absent in each genome assembly if a box is white (unshaded). Bolded and underlined items in the ‘GENE’ column represent genes that are meiosis-specific. The ‘FUNCTIONAL ROLE’ column describes the meiotic function of each gene.

*Transposable element discovery:* We annotated TEs in the *P. antipodarum* genome using a variety of approaches (see Supplementary Information for details). In brief, RepeatModeler (using RepeatScout/RECON) found 2294 repetitive element families and the LTRPipeline found an additional 253 families, 122 of which were removed in clustering for redundant sequences, for a total of 2425 repetitive element families. We curated these repetitive element families with the automated pipeline MCHelper, resulting in 957 classified TE families (Supplementary Table S1, and see Supplementary Information for description of TE classifications).

### Evidence for a recent WGD and post-WGD evolution in *P. antipodarum* genome

*Allele sequencing depth at heterozygous sites:* Assuming unbiased sequencing, heterozygous sites are expected to exhibit approximately equal depth of coverage in a diploid (i.e., A:B). By contrast, heterozygous sites in triploids are expected to reflect a 2:1 or 1:2 coverage ratio (i.e., AA:B; A:BB), while tetraploids could in principle display the full range of 1:3, 1:1, and 3:1 coverage ratios (i.e., AAA:B; AA:BB; A:BBB). This expectation allowed for a straightforward test of diploidy using the relative depth of coverage for each allele at heterozygous sites, revealing substantial evidence of non-equal coverage across the *P. antipodarum* genome (Figure 3A). While many sites across the genome fit the null diploid expectation (i.e., 362125/3438187, ∼10.5% of heterozygous sites exhibited minor allele frequencies between 0.45 – 0.5), a large fraction of the genome nevertheless exhibited the 1:2 (2:1) coverage pattern (378587/3438187, ∼11.0% of heterozygous sites exhibited minor allele frequencies between 0.305 – 0.355) or the 1:3 (3:1) coverage pattern (435622/3438187, ∼12.7% of heterozygous sites exhibited minor allele frequencies between 0.225 – 0.275). This distribution of sequencing coverage is not compatible with pure diploidy but indicates a more complex genome structure.

**Figure 3.**
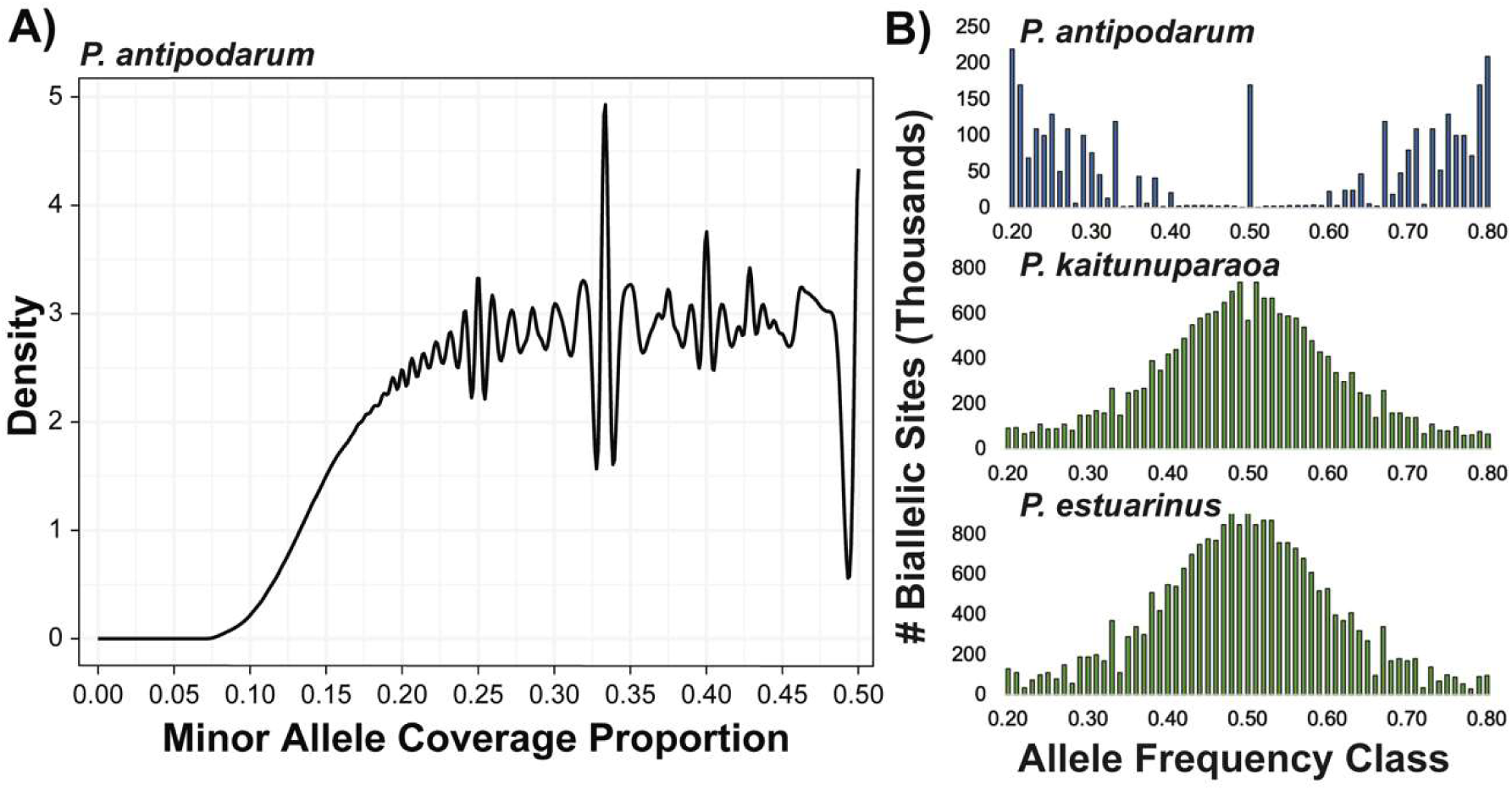
Allelic depth and site-frequency spectra do not meet expectations for diploidy in *P. antipodarum*. A) Density plot summarizing relative coverage proportion of the minor allele at heterozygous positions, identified in the collapsed assembly. The notable peaks at 0.25 (tetraploidy) and 0.33 (triploidy) indicate that the *P. antipodarum* genome does not fit the typical expectations of diploidy. B) Whole-genome site-frequency spectra for *top* – *P. antipodarum* reads (Alex Yellow) mapped to the uncollapsed *P. antipodarum* assembly, *middle* – *P. estuarinus* reads mapped to *P. estuarinus* genome, and *bottom* – *P. kaitunuparaoa* reads mapped to *P. kaitunuparaoa* genome.

*Site-frequency spectrum analysis:* We next estimated genome copy number *in silico* using nQuire [49], leveraging the same data as for the allele depth analysis. The collapsed *P*. *antipodarum* assembly contained 1.59 M high quality biallelic sites (0.16% of the genome assembly), the *P. estuarinus* assembly contained 11.50 M biallelic sites (2.23% of the assembly), and the *P. kaitunuparaoa* assembly contained 9.32 M biallelic sites (1.56% of the assembly) (Figure 3B). Alleles at biallelic sites are expected to exhibit a unimodal frequency distribution centered at 0.5 in diploids, while triploids are expected to have peaks at 0.33 and 0.66, and tetraploids are expected to have a trimodal allele frequency distribution with peaks at 0.25, 0.5, and 0.75. We evaluated these distributions using log-likelihoods (1/*Δ*log*L*) for each model comparison, whereby the larger the number, the better the model fits the data. Diploid relatives *P. estuarinus* (1/*Δ*log*L*: diploid = 5.0; triploid = 0.10; tetraploid=0.01) and *P. kaitunuparaoa* (1/*Δ*log*L*: diploid = 5.0; triploid = 0.14; tetraploid = 0.02) both displayed SFS that best fit a diploid model, whereas *P. antipodarum* (1/*Δ*log*L*: diploid = 0.53; triploid = 0.91; tetraploid = 10.0) had a SFS that best fit a tetraploid model (Figure 3B). The unexpectedly complex distribution observed in the *P*. *antipodarum* assembly prompted us to estimate the SFS in each of the 100 scaffolds with the highest number of biallelic sites. Of these, 16 scaffolds best fit a diploid model, two best fit a triploid model, and 81 best fit a tetraploid model (Supplementary Figure S4; Supplementary Table S2). One contig (tig00149948) was an equally good fit for the triploid and tetraploid models (1/*Δ*log*L*: diploid = 0.003; 1/*Δ*log*L*: triploid = 0.01; tetraploid = 0.01). By contrast, all 100 scaffolds with the highest number of biallelic sites in both the *P. estuarinus* and *P. kaitunuparaoa* assemblies best fit a diploid model (Supplementary Tables S3, S4).

*Duplicate gene content and evolutionary history:* One of the hallmarks of WGDs (*vs.*, for example, repeat expansion or structural rearrangements) is instantaneous doubling of gene content across the genome. We first evaluated whether any of the three *Potamopyrgus* species fit this expectation of WGD by using Orthofinder to infer genome-wide homology in the *P. antipodarum* (uncollapsed), *P. estuarinus*, and *P. kaitunuparaoa* proteomes. The OrthoFinder run produced 25,380 homologous groups containing >1 gene (orthogroups, Supplementary Figure S5), 7837 (30.9%) of which were single-copy in all three species (Figure 4A). The next most prevalent pattern among orthogroups was of genes that were double copy in *P. antipodarum* but single copy in the other two species (n = 3040, 12.0%). This group was more than six times larger than the number of analogous orthogroups in which two gene copies were represented by *P. kaitunuparaoa* (487 orthogroups) or by *P. estuarinus* (442 orthogroups) but only a single copy from each of the other two species (FET Odds Ratio: 6.96; 95% CI: 6.31 – 7.69; *p* < 0.0001).

**Figure 4.**
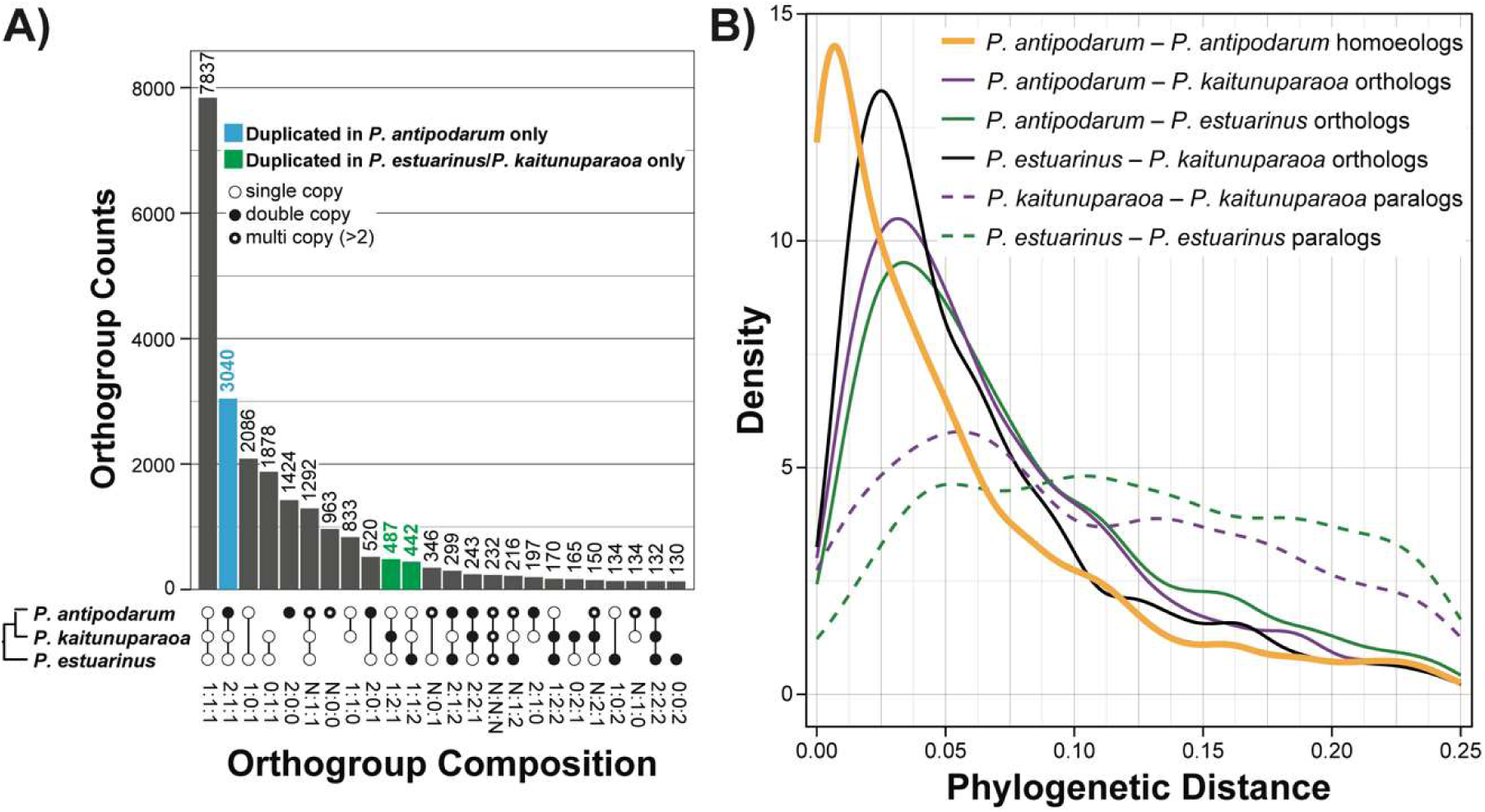
Duplicate gene content and phylogenetic evidence of whole-genome duplication. A) Upset plot describing orthologous gene group composition in *P. antipodarum*, *P. kaitunuparaoa*, and *P. estuarinus*. *Top –* Histogram of orthogroup number within bins. The blue column represents the 2:1:1 orthogroup bin, for which *P. antipodarum* has two gene copies, while *P. kaitunuparaoa* and *P. estuarinus* have only a single copy. Green columns represent analogous orthogroups in which *P. kaitunuparaoa* and *P. estuarinus* have double-copy genes, but the other species only have a single copy. *Bottom* – Orthogroup bins describing gene content in which white circles represent single-copy genes, black circles represent double-copy genes, and black circles encircling white circles represent multi-copy (i.e., >2) genes. Absence of a circle indicates the absence of a gene from that particular species. The full distribution is available in Supplementary Figure S5. B) Density plots of pairwise phylogenetic distance (i.e., patristic distance) from quartet trees (i.e., 2:1:1; 1:2:1; and 1:1:2 orthogroup bins). The solid orange line refers to phylogenetic distance between gene duplicates specific to *P. antipodarum*. The solid purple line represents the mean phylogenetic distance of both *P. antipodarum* gene copies to *P. kaitunuparaoa.* The solid green line represents the mean phylogenetic distance of both *P. antipodarum* gene copies to *P. estuarinus.* The dashed purple line represents phylogenetic distance between *P. kaitunuparaoa*-specific gene duplicates (1:2:1 orthogroup bin). The dashed green line represents phylogenetic distance between *P. estuarinus*-specific gene duplicates (1:1:2 orthogroup bin).

We inferred gene trees for both the triplet orthogroups (i.e., 1:1:1) and all possible quartets (i.e., 2:1:1; 1:2:1; and 1:2:2) to estimate phylogenetic distance between gene duplicates. These gene trees (Supplementary Figure S6) revealed two patterns: 1) duplicate genes in *P. antipodarum* are much more closely related to each other than either are to *P. kaitunuparaoa* or to *P. estuarinus* (Table 1, Figure 4B, Supplementary Table S5), and 2) all pairs of duplicate genes in *P. antipodarum* appear to have originated around the same time (peak divergence = 0.0070 substitutions/site; Figure 4B). Neither pattern applies to *P. kaitunuparaoa* or *P. estuarinus* paralogs. This finding indicates that the gene duplicates (hereafter homoeologs) found in the *P. antipodarum* uncollapsed contigs appear to have originated from a WGD, whereas the paralogs present in the other two species reflect the background rate of individual gene duplication.

**Table 1.** Gene tree summary statistics.

| Orthogroup bin | Median Paralog Divergence (95% CI)<sup>a</sup> | Median *P. antipodarum* - *P. kaitunuparaoa* Divergence (95% CI) | Median *P. antipodarum* - *P. estuarinus* Divergence (95% CI) | Median *P. kaitunuparaoa* - *P. estuarinus* Divergence (95% CI) |
| --- | --- | --- | --- | --- |
| *Double-copy genes* |  |  |  |  |
| 2:1:1<sup>b</sup><br>(n = 3040) | 0.038<br>(0.000 – 0.522) | 0.062<br>(0.007 – 0.688) | 0.063<br>(0.008 – 0.608) | 0.065<br>(0.008 – 0.671) |
| 1:2:1<sup>c</sup><br>(n = 487) | 0.219<br>(0.000 – 34.58) | 0.173<br>(0.012 – 17.33) | 0.073<br>(0.010 – 1.82) | 0.169<br>(0.016 – 17.33) |
| 1:1:2<sup>d</sup><br>(n = 442) | 0.312<br>(0.022 – 34.54) | 0.067<br>(0.008 – 1.18) | 0.216<br>(0.027 – 17.28) | 0.216<br>(0.027 – 17.28) |
| **Concatenated**<br>**2:1:1<sup>e</sup>** | **0.054** | **0.072** | **0.079** | **0.063** |
| *Single-copy genes* |  |  |  |  |
| 1:1:1<sup>f</sup><br>(n = 7837) | N/A | 0.040<br>(0.004 – 0.688) | 0.046<br>(0.006 – 0.669) | 0.040<br>(0.006 – 0.541) |
| **Concatenated**<br>**1:1:1<sup>g</sup>** | **N/A** | **0.053**<br>**(0.052 – 0.054)** | **0.057**<br>**(0.057 – 0.058)** | **0.052**<br>**(0.052 – 0.053)** |

<sup>a</sup> – 95% confidence intervals were determined from the value of the 2.5<sup>th</sup> percentile and 97.5<sup>th</sup> percentile of patristic distances from all trees present in the orthogroup bin.
<sup>b</sup> – All orthogroups in this bin contain two *P. antipodarum* sequences and one sequence from each of the other two species. Paralog divergence here refers to the total patristic distance from one *P. antipodarum* copy to the other. Divergence from the other species was estimated as the mean between the two branches for each comparison.
<sup>c</sup> – All orthogroups in this bin contain two *P. kaitunuparaoa* sequences and one sequence from each of the other two species. Paralog divergence here refers to the total patristic distance from one *P. kaitunuparaoa* copy to the other. Divergence from the other species was estimated as the mean between the two branches for each comparison.
<sup>d</sup> – All orthogroups in this bin contain two *P. estuarinus* sequences and one sequence from each of the other two species. Paralog divergence here refers to the total patristic distance from one *P. estuarinus* copy to the other. Divergence from the other species was estimated as the mean between the two branches for each comparison.
<sup>e</sup> – All 2:1:1 orthogroups were concatenated in this phylogenetic analysis. Variance estimates are not available from bootstrap replicates produced by RAxML.
<sup>f</sup> – All orthogroups in this bin contain a single sequence from each species. Because only a single unrooted tree topology is possible for a tree with three taxa, only branch lengths were estimated using PhyML in this analysis.
<sup>g</sup> – All 1:1:1 orthogroups were concatenated in this phylogenetic analysis. 95% CI estimates of patristic distances are from 100 bootstrap replicates in PhyML.

*Conservation of synteny of gene duplicates:* Another hallmark of WGD and a useful tool for determining when these genome duplication events occurred is the extent of conserved gene collinearity (*i.e*., synteny) across groups of duplicated genes arrayed along a chromosome [53]. The potential for extensive recombination-derived structural rearrangements in the wake of WGD [54,55] can make this pattern challenging to identify, but even genome-wide blocks of microsynteny (*i.e.*, 10-50 genes) can provide powerful evidence of homoeology [56]. Here, we leveraged the collapsed *P. antipodarum* scaffolds to serve as anchor points with which to compare synteny in the uncollapsed contigs, with *P. kaitunuparaoa* and *P. estuarinus* as outgroup references. We found evidence of global microsynteny across *P. antipodarum* homoeologs, interrupted by complex structural rearrangements (Supplementary Figure S7). As an exemplar, we performed an in-depth characterization of microsynteny on Scaffold 14 from the collapsed assembly (Figure 5), as it featured the longest block of 2:1:1 orthogroups along a single scaffold (n = 57). In this 3-Mbp genomic region, we can simultaneously observe the maintenance of synteny among homoeologous gene pairs, individual gene loss, and even structural variation across subgenomes. There was perfect microsynteny across the genus for 53/57 (93%) of 2:1:1 orthogroups, whereas only 45/62 (73%) of single-copy orthogroups exhibited conserved microsynteny with the other *Potamopyrgus* species. The greater degree of structural rearrangement among single-copy genes potentially reflects the mechanism of gene loss (*i.e.*, via ectopic recombination). Overall, these synteny data provide clear evidence that a WGD event followed by complex and extensive post-WGD chromosomal rearrangements underpin genome evolution in *P. antipodarum*.

**Figure 5.**
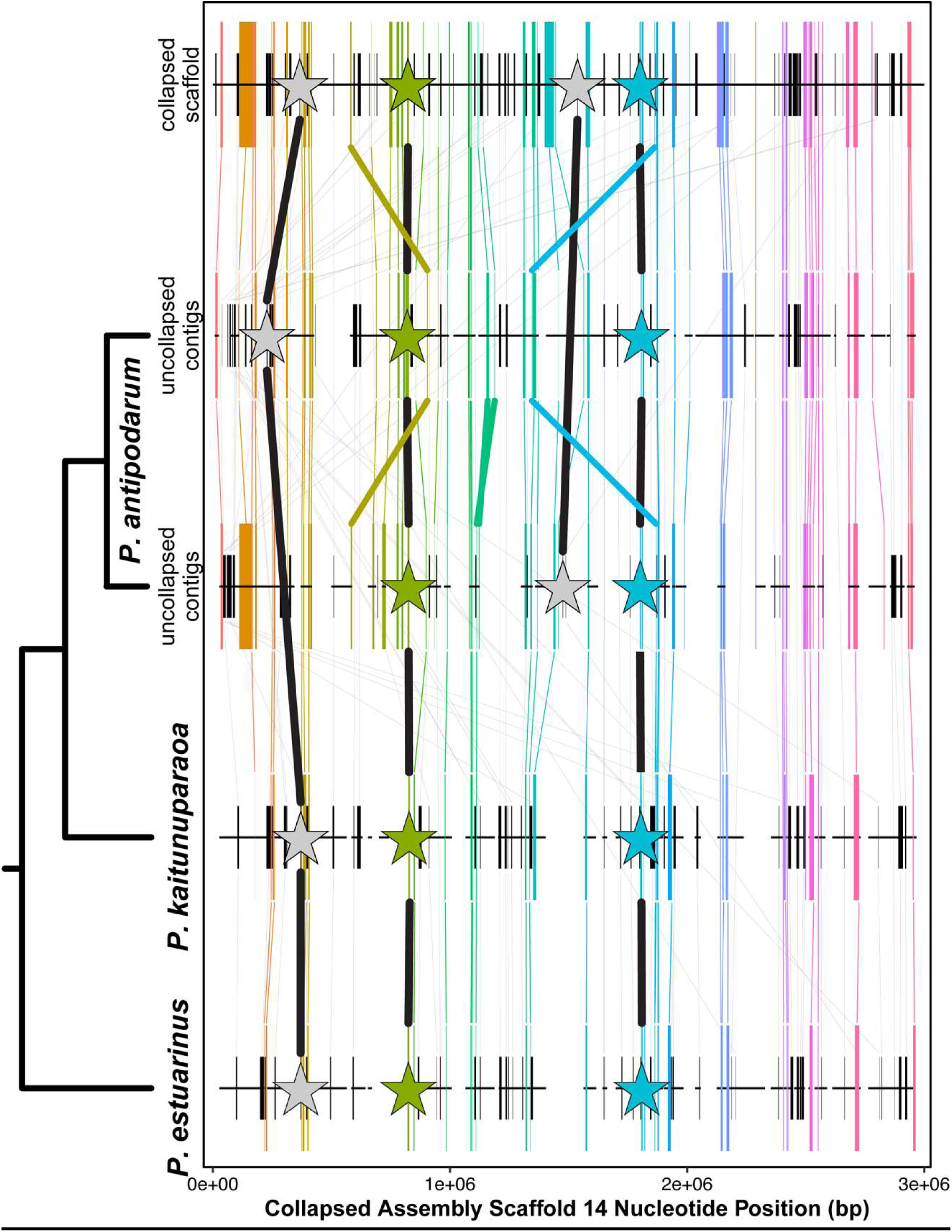
Microsynteny of duplicated and single-copy genes in the *P. antipodarum* genome. Orthologous genes mapped along Scaffold 14 of the collapsed assembly, along with their corresponding positions on the uncollapsed contigs and in the *P. kaitunuparaoa* and *P. estuarinus* assemblies. Genes that were double copy in *P. antipodarum* uncollapsed contigs, but single copy in all other assemblies (i.e., 2:1:1 orthogroup bins, n = 57) are represented by tall colored boxes, with the width reflecting the length of the gene. Genes that are single copy in all species (i.e., 1:1:1 orthogroup bins, n = 62) are represented by short black boxes, with the width of the box representing gene length. Lines connecting orthologs are colored for 2:1:1 orthogroups and grey for 1:1:1 orthogroups. Large stars represent BUSCO genes (connected by black lines), with colored stars being 2:1:1 genes and grey stars being 1:1:1 genes. Thick colored lines reflect rearrangements found in 2:1:1 genes.

*Transposable element abundance does not explain the increase in P. antipodarum genome size:* To rule out the possibility that TE expansion is responsible for the larger genome size of *P. antipodarum*, we compared TE proportion and total genomic TE content across three *Potamopyrgus* species. DNApipeTE estimates of genomic TE content indicated that *P. antipodarum* harbors more TEs than its congeners, with 42% of genomic reads mapping back to putative TEs *vs*. ∼25% for *P. estuarinus* and *P. kaitunuparaoa* (Supplementary Table S6). The RepeatMasker results are overtly similar, with TEs representing 45.6% of the *P. antipodarum* genome and only 26.1% of the *P. estuarinus* genome (Table 2). The outcome of the analysis of the masked and collapsed version of the *P. antipodarum* genome was virtually identical, with 42.8% TEs (Figure 6A). The congruencies between our assembly-based and read-based measures of TE abundance, including the two levels of assembly for the *P. antipodarum* genome assembly, indicate that we accurately captured the proportional differences in TE load across *Potamopyrgus* species.

**Figure 6.**
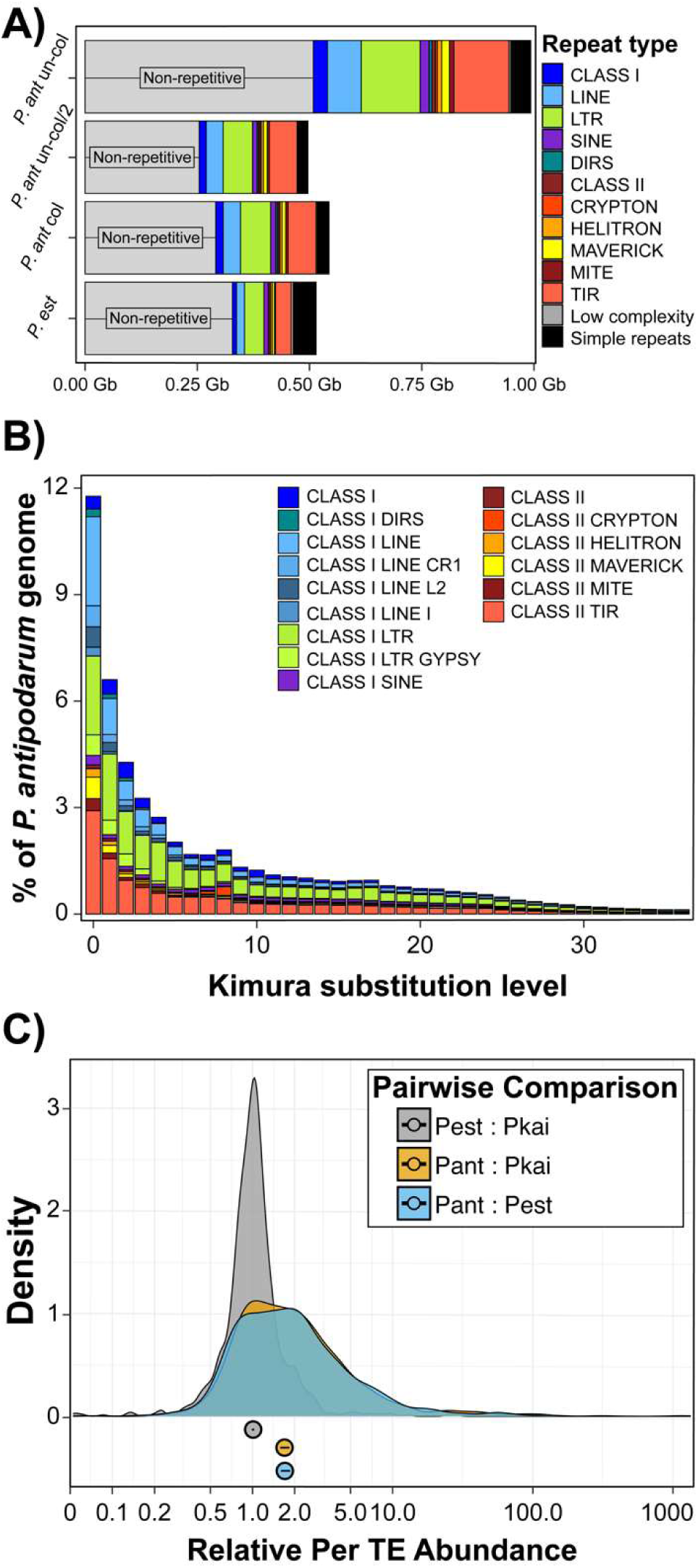
Transposable element expansion in the *P. antipodarum* genome. A) Comparison of repetitive sequence content across *P. antipodarum* genome assembly versions and *P. estuarinus* based on RepeatMasker results using the curated *P. antipodarum* transposable element (TE) library. Genome size for each assembly is reflected in the X-axis. Repeat type (classes and major divisions of TE types) are color coded, along with “Non-repetitive” genomic content (i.e., not masked by any repeat category). Genome assemblies are as such, “*P. ant* un-col”: uncollapsed (contigs) genome assembly for *P. antipodarum*; “*P. ant* un-col/2”: results for the uncollapsed genome assembly for *P. antipodarum* divided by 2; “*P. ant* col”: collapsed (scaffold) assembly for *P. antipodarum*; “*P. est*”: *P. estuarinus* genome assembly. B) Repeat landscape for *P. antipodarum*, depicting past TE activity in the genome. X axis: Kimura substitution level relative to the family consensus sequence, reflective of TE family age. Younger TEs (i.e., more recently inserted into the genome) have lower levels of pairwise divergence compared to older TEs. Y axis: percent of the genome made up by each TE group. C) Relative abundance of TEs in pairwise species comparisons. Copy numbers estimated from read mapping depth via DeviaTE for 942 TE families, shown here as ratios of copy numbers between species, *e.g.*, for a “relative copy number” of 1.0, the two species in comparison have identical copy number. Three pairwise comparisons, *e.g.*, *P. antipodarum*: *P. estuarinus* (Pant: Pest) and *P. antipodarum: P. kaitunuparaoa* (Pant: Pkai) are shown as density plots of 942 relative values, with median and 95% CI shown below. Example highlighted in text, DNA transposon rnd-5_family-12008, copy numbers: Pkai = 4.306, Pest = 6.943, Pant = 197.091; relative values: Pest:Pkai = 1.6, Pant:Pkai = 45.8, Pant:Pest = 28.4.

**Table 2.** Transposable elements in *P. antipodarum* compared to other *Potamopyrgus* species.

| Repeat classification |  | DNApipeTE |  |  | RepeatMasker |  |
| --- | --- | --- | --- | --- | --- | --- |
|  |  | *P. antipodarum* | *P. estuarinus* | *P. kaitunuparaoa* | *P. antipodarum* | *P. estuarinus* |
| Class I |  |  |  |  |  |  |
|  | LTR | 2.54% | 0.01% | 1.68% | 13.23% | 8.43% |
|  | LINE | 4.90% | 2.28% | 1.91% | 7.58% | 3.42% |
|  | SINE | 0.40% | 0.77% | 0.37% | 2.02% | 1.85% |
|  | DIRS | n/a | n/a | n/a | 0.64% | 0.11% |
|  | other | n/a | n/a | n/a | 2.52% | 1.71% |
| Class II |  |  |  |  |  |  |
|  | TIR | 3.66% | 3.19% | 3.02% | 13.25% | 7.12% |
|  | Maverick | n/a | n/a | n/a | 1.72% | 1.04% |
|  | Helitron | 0.36% | 0.21% | 0.33% | 1.01% | 0.73% |
|  | Crypton | n/a | n/a | n/a | 0.43% | 0.08% |
|  | other | n/a | n/a | n/a | 0.75% | 0.77% |
| Other repeats |  |  |  |  |  |  |
|  | Low complexity | 0.20% | 0.48% | 0.56% | 0.42% | 0.94% |
|  | Simple | 2.26% | 3.89% | 4.22% | 4.45% | 9.86% |
|  | other | 0.34% | 0.95% | 1.04% | n/a | n/a |
| Unclassified |  | 30.16% | 17.82% | 17.24% | n/a | n/a |
| **Total TE** |  | **42.02%** | **25.75%** | **24.56%** | **43.15%** | **25.26%** |
| **Total repeat** |  | **44.83%** | **31.06%** | **30.37%** | **48.02%** | **36.06%** |

Although our results revealed a considerably higher burden of TEs in *P. antipodarum* relative to two congeneric species, the genome size disparities between *P. antipodarum* and *P. estuarinus* cannot be explained by TEs alone. Notably, the total bp of TEs in the *P. antipodarum* genome assembly was about 3.3x higher than the *P. estuarinus* assembly, while the non-repetitive portion of the *P. antipodarum* genome was still about 1.5x greater than in *P. estuarinus*, closely matching the 1.8x more predicted genes.

*Recent “bursts” of TE activity in* P. antipodarum *genome:* Over 20% of the *P. antipodarum* genome is made up of TEs with within-family divergences of 2% or less (Figure 6B). The proportion of TE sequence belonging to the 0-2% divergence bins was more than double the proportion of TEs sequence in the 3-5% divergence bins, demonstrating a global increase in TEs across all classes. Read-mapping approaches also revealed higher abundances of TEs (on a per-element basis) in *P. antipodarum* relative to *P. estuarinus* and *P. kaitunuparaoa* (Figure 6C). While some elements demonstrated no notable difference in copy number across species (see example in Supplementary Information), these comparative results demonstrate a systematic increase in TE abundance in *P. antipodarum* relative to its close relatives, and in one case, a likely active DIRS retrotransposon exclusively found in *P. antipodarum* (Supplementary Figure S9, and see supplementary text for details).

We next asked whether there existed dramatic differences in copy number and relative abundance of particular TE families across species as an indicator of “bursts” of TE activity. For this analysis, we applied an arbitrary threshold of differences in copy number of at least 100 and at least 5x difference in relative abundance. For *P. antipodarum vs*. *P. estuarinus*, 24 TEs met these criteria, while 27 TEs met the criteria for *P. antipodarum vs*. *P. kaitunuparaoa* (with 20 in common across these two pairs). There were no other comparisons that reached these thresholds across any of the other pairwise species combinations, with the exception of one TE family in *P. estuarinus* compared to *P. kaitunuparaoa* (110.4 copies in *P. antipodarum*, 284.2 copies in *P. estuarinus*, and 28.7 copies in *P. kaitunuparaoa*). This result demonstrates that recent accumulation of TEs is characteristic only in *P. antipodarum*.

We also examined the RepeatMasker results to identify elements with likely ongoing activity to ask whether these elements are more abundant in *P. antipodarum*. The criteria we considered were the sequence divergence between copies of TEs of the same family and “completeness” of copies in the *P. antipodarum* genome assembly (i.e., how long is the average copy length compared to the reference length of the TE?). The element with the lowest Kimura substitution level (0.3%) was a *Mariner* DNA transposon that also exhibits evidence of activity burst in *P. antipodarum* compared to its congeners, with nearly 200 more copies (Supplementary Figure S10, and see Supplementary Information for additional details). The TE with the longest average genomic insertion (8,024 bp *vs*. 13,140 bp reference length) was classified as a *Maverick* DNA transposon that also fit out criteria for burst. This *Maverick* element accounts for about 3.5 Mb of genomic sequence in *P. antipodarum vs*. about 0.05 Mb in *P. antipodarum* (see Supplementary Information for additional details).

## Discussion

We produced the first high-quality reference genome assembly for *Potamopyrgus antipodarum*, a New Zealand freshwater snail that has risen to prominence as a model system for the study of sexual reproduction, host-parasite coevolution, genome size variation, and invasion biology. Analysis of this genome assembly also revealed multiple strong lines of evidence for an unexpected and relatively recent whole-genome duplication (WGD) unique to *P. antipodarum* and rampant post-WGD activation of TEs.

Together, our analyses point to a scenario where the differential retention and loss of extra chromosome copies post-WGD has resulted in a mostly tetraploid mosaic genome with a substantial minority of triploid and diploid genomic regions. This complexity is consistent with expectations of rediploidization following a WGD [54,55]. The extent to which the redundant genome copies are retained or expunged via random genetic drift or natural selection is an exciting open question. Both selection [57] and genetic drift [58] have been implicated in the (non)random genomic winnowing that generally occurs after WGD as rediploidization progresses. The variable nuclear genome size even with respect to diploid sexual *P. antipodarum* [38,51] means that these questions can be addressed with comparisons between *P. antipodarum* that differ in nuclear DNA content. Such analyses will illuminate how selection and genetic drift operate under conditions where excess genomic material has been gained [59,60].

### General Implications of WGD

We unexpectedly discovered multiple strong lines of evidence for a very recent WGD in the ancestor of extant *P. antipodarum*. WGD events are among the most profound mutational changes observed in nature (reviewed in [61]), with effects ranging from the genome (*e.g.*, [54,62,63]) and cell (*e.g.*, [64–66]) to the organism (*e.g.*, [67–69]) and population (*e.g.*, [70–74]). The genome-wide redundancy that results from WGDs has long been thought to provide the fuel for evolutionary innovation [63,75], with important adaptations like seeds [76], flowers [77], improvements in visual systems [78], and many other morphological and developmental innovations [79–82] owing their origins to WGDs.

Among the most perplexing consequences of WGDs is the seemingly inevitable return to diploidy following genome duplication [83]. Indeed, although the vast majority of extant eukaryotes are diploid, almost all have experienced WGD events in their evolutionary past [84,85], including humans [53]. The rate and pattern of rediploidization is therefore of great interest to evolutionary biologists, and much has been learned over recent years, especially with the explosion of plant genomic resources [72,86,87]. Whether these lessons extend to animals is unclear, in large part because only two animal systems (i.e., *Xenopus* [88] and salmonids [89,90]) have the necessary genomic resources. Comparing a diverse set of independent animal WGDs is therefore critical to identifying the “rules” (if any exist) of post-polyploid genome evolution. Our *P. antipodarum* genome assembly provides a valuable advance in this respect [34,91].

### Meiosis genes in the context of WGD

In the *P*. *antipodarum* genome assembly, 38/44 (86%) of the meiosis genes queried are present in at least one copy, and at least one of those copies is intact. The presence and likely intact function of most of the genes needed to maintain meiosis [52] in the inbred sexual *P. antipodarum* line that we sequenced here as well as the nearly identical meiosis gene complement in closely related obligately sexual species is as expected if purifying selection is maintaining gene function required for obligate sex. Some of these duplicated genes have disrupted reading frames or are missing portions of exons, suggestive of rediploidization following the WGD [53,85,92,93]. Nevertheless, that all genes present in *P*. *antipodarum* are also present in closely related congeners as well as in more distantly related taxa provides solid evidence that these genes are indeed maintained in *P. antipodarum*.

Conspicuous meiosis gene absences included RECQ3, TIM2, and CycA. Because these genes were absent, their absence might characterize the subclass Caenogastropoda. The absence of meiosis genes from obligately sexual species is not unprecedented [52,94]. Indeed, these “incomplete” meiotic toolkits seem to be the rule rather than the exception, found in taxa from diatoms [95] and choanoflagellates [96] to *Trichomonas [97]* and many more [98–101].

### Transposable element proliferation following WGD

Among the expected consequences of WGD are altered regulation and evolution of transposable elements [102,103]. In particular, polyploidization has long been thought of as a “genomic shock” that enables bursts of TE activity [104]. While polyploids tend to have more TEs than diploid relatives, the mechanisms underlying this phenomenon remains an open question [102,105], particularly concerning the relative contributions of TE activity and efficacy of natural selection to remove TE insertions in polyploids [106–108]. Our analysis of the *P. antipodarum* genome revealed a global increase in TEs relative to its close relatives and evidence of recent bursts of multiple specific TE families.

Transposable elements are important contributors to genome size variation [109] and may account for dramatic differences in genome size between even closely related species (*e.g.*, [110–112]), with expansion of a single TE family capable of rapidly increasing genome size [113]. Although our results revealed a considerably higher burden of TEs in *P. antipodarum* relative to congeners (*e.g.*, 3.3x more total DNA in genome assemblies), the higher amount (1.5-1.8x) of DNA in genes and in non-repetitive genomic regions means that nuclear genome size disparities between *P. antipodarum* and *P. estuarinus* cannot be explained by TEs alone. It is possible that the genic region of the *P. antipodarum* genome is rediploidizing at a higher rate than repetitive regions, which may account for the asymmetry in expanded DNA regions relative to *P. estuarinus*. Post-WGD TE proliferation may also rapidly add substantial quantities of DNA to the genome, driving up the proportional differences in TE abundance. As an example from *P. antipodarum*, we identified a 13 Kb *Maverick* DNA transposon with hundreds of more copies in the *P. antipodarum* genome compared to its congeners that alone accounts for a 3.5 Mb difference in genomic sequence.

Along with this *Maverick* transposon, we identified several other likely active elements representing different TE classes that exhibit signs of activity bursts in *P. antipodarum*, *e.g.*, low sequence divergence across – mostly – complete copies and dramatically greater abundance than in other *Potamopyrgus* species. Population-level and polymorphism data from multiple species of *Potamopyrgus* will allow for further tests of the influence of WGD on TE evolution and for evaluating whether the WGD might have increased TE activity in *P. antipodarum*. Finally, the apparently recent invasion of some TEs raises important questions about genome stability and susceptibility to TE colonization and mobilization and will allow us to use *P. antipodarum* to explore the influence of asexuality on TE evolution, including direct evaluation of the conflicting expectations regarding how the absence of sex will affect TE proliferation [114–123].

### Could the WGD underpin the high levels of variation and the evolution of asexual reproduction observed in *P. antipodarum?*

*Potamopyrgus antipodarum* is notable as one of the few natural systems with coexisting and competing obligately sexual and asexual forms [9]. While some other animal taxa do experience occasional transitions to asexual reproduction [124,125], most of these cases are tied to hybridization and are observed only rarely [126], *vs*. the evidence for many and frequent such transitions in *P. antipodarum* [15,127,128]. *P. antipodarum* is the only species in the large Tateidae family known to have evolved asexual reproduction [129], and there are only two other gastropod taxa for which parthenogenesis has been documented (reviewed in [130]). These snails harbor high phenotypic diversity relative to other New Zealand congeners [131–134], and asexual assemblages of *P. antipodarum* are rich in genetic diversity relative to other naturally occurring asexual populations [127,128]. This snail also inhabits a remarkably wide range of habitats [131,134] and has established invasive populations across the world [31].

Could a recent WGD help to explain why *P. antipodarum* is so unique? There is a growing body of evidence that WGD can catalyze the evolution of novel phenotypes (recently reviewed in [61]). We are unaware of studies demonstrating a causal link between WGD and frequent transitions to parthenogenesis, but there are several cases of strong links between WGD and other innovations in key features of reproductive biology (*e.g.*, [79,135]). The extra copies of chromosomes and genes that characterize WGD can also provide a means of compensating for some of the hypothesized negative consequences of asexual reproduction (*e.g.*, mutation accumulation, lack of genetic variability) *via* mutational masking [124,136] and as a new source of genetic variation (*e.g.*, [137,138]). Moreover, the rediploidization that follows WGD could lead to broad-scale genomic incompatibilities resulting from lineage-specific genome fractionation, from which the only evolutionary escape is asexuality. Thus, WGD may provide a scenario in which asexuality is able to gain a foothold in natural populations. The resources we present here provide a powerful means forward to begin targeted evaluation of whether and how WGD might enable the repeated separate transitions to obligate asexuality that make *P. antipodarum* so remarkable.

## Summary & Conclusions

The *P. antipodarum* genome assembly that we have produced represents a unique resource that will be of use to the many scientists interested in a host of important open questions in biology, from the maintenance of sex and the evolution of biological invasions to the consequences of WGD and the genomic drivers of host-parasite coevolution. Our genome assembly will enable downstream work targeting the genomic or genetic regions involved in the frequent transitions to asexual reproduction in *P. antipodarum*. We have also brought together multiple lines of evidence, including some new ways to evaluate genomes for the presence of WGD, to demonstrate a recent WGD in *P. antipodarum* that appears to be “caught in the act” of the process of rediploidization. This discovery opens the door to using different *P. antipodarum* lineages to evaluate the extent to which genomic incompatibilities produced by rediploidization proceeding independently in allopatric lineages could contribute to frequent transitions to asexuality in this system.

## DATA AVAILABILITY

All raw sequencing data are available at NCBI in the SRA (SRR29047464, SRR29047466, SRR29099380, SRR29099381, SRR29099382, SRR29 099389, SRR32688197) under BioProject PRJNA717745. The uncollapsed contig assembly is currently being processed at NCBI, while both the uncollapsed and collapsed assemblies, transcriptome assemblies, all annotations, and gene trees are available on Zenodo (https://doi.org/10.5281/zenodo.15015201). All scripts developed for this project, as well as all supplemental information and flow cytometry data are available on GitHub (https://github.com/jsharbrough/potamomics). Alex Yellow snails are available on request.

## Acknowledgments

We thank Curt Lively for providing us with members of the Alex Yellow culture. We thank Katelyn Larkin for project support, and especially DNA extraction optimization. We acknowledge Marissa Roseman, Ben Ripperger, and Mohammed Farooqi for snail maintenance, Gery Hehman for RNA extraction, library preparation, and sequencing, and DNA library preparation and sequencing, Dovetail Genomics for high-molecular weight DNA extractions and Chicago library construction and sequencing, Arizona Genomics Institute for PacBio DNA sequencing, Matthew Brockman for assistance with computational resources, Emily Jalinsky for snail illustrations, and Mary Morgan-Richards for advice regarding appropriate credit for acknowledgment and guardianship. The authors acknowledge the kaitiaki (guardians) of the endemic species, the people who are the traditional landholders of Takamana, where the pūpū were collected: Arowhenua.

We acknowledge funding from NSF-MCB 1122176 (Neiman, Logsdon, Boore); NSF-DEB 1753851 (Neiman); NSF-DEB 1310825 (Neiman, Sharbrough); NSF-DEB 1601242 (Neiman, McElroy); Carver Biomedical Trust 18-5081 (Neiman); Iowa Office for Undergraduate Research Funding (Neiman); Iowa Science Foundation (Neiman); NSF DEB-175433 (McElroy); NSF DEB-2148203 (McElroy); NSF OAC-2322260 (Sharbrough); Colorado State University (Sharbrough); New Mexico Institute of Mining and Technology (Sharbrough); New Mexico INBRE (Sharbrough). This work also used the Alpine high-performance computing resource at the University of Colorado Boulder. Alpine is jointly funded by the University of Colorado-Boulder, the University of Colorado-Anschutz, and Colorado State University and with support from NSF grants OAC-2201538 and OAC-2322260.

