## Supplementary material for "Whole-genome sequence of *Potamopyrgus antipodarum*—a model system for the maintenance of sexual reproduction—reveals a recent whole-genome duplication": Supplementary Information.pdf

### Methods

##### Genomic Sequencing

Genomic sequence data from Alex Yellow was generated with Illumina HiSeq and PacBio CLR sequencing platforms, as well as a Dovetail Chicago *in vitro* proximity-ligated library [1] that was sequenced on an Illumina HiSeq. All snails used for sequencing were housed in standard laboratory conditions (16ºC constant-temperature room, 16:8 hour light/dark cycle, fed dried *Spirulina* 3x/week; following ref. [2]). All Alex Yellow snails used for DNA extraction for these sequencing runs were non-brooding (i.e., did not contain eggs or embryos) adult females. When we did DNA extractions ourselves, we used the Zymo Clean and Concentrate Kit (Zymo, Irvine, USA) to clean the DNA extractions and then resuspended the DNA in 30-100 µL of T-low-E buffer (10mM Tris pH 8.0, 0.1mM EDTA) prior to library preparation. Each sample was analyzed on a NanoDrop 1000 to determine DNA concentration and purity. We only proceeded with DNA extractions that contained 200-1000 ng DNA and for which the 260/280 was ~1.8. 100 ng of each sample was run on a 0.8% agarose gel to confirm the integrity of the sample as represented by the presence of a single band of high-molecular weight DNA and to identify visible contaminants. All samples that we used for sequencing showed a single strong nucleic acid band on this gel.

*Illumina HiSeq (2014):* We extracted DNA following a modified CHAOS protocol [3,4] from 15 Alex Yellow individuals and then pooled the 15 individual DNA extractions for Illumina TruSeq DNA PCR-free library preparation. We sequenced this library in a single lane of 2x150 paired-end sequencing on an Illumina Hi-Seq 2500. SRA Accession: SRR29047466.

*Dovetail Proximity-Ligated Library (2017):* We sent the tissue of a single Alex Yellow female for high-molecular-weight (HMW) DNA extraction to Dovetail Genomics (Santa Cruz, USA), followed by proximity ligation (Chicago) library preparation and sequencing. Long-ranged mate pairs were produced using the Chicago method [1] and sequenced by Dovetail Genomics (Santa Cruz, USA). Briefly, 500 ng of HMW genomic DNA was reconstituted into chromatin *in vitro* and fixed with formaldehyde. Fixed chromatin was digested with MboI, the 5′ overhangs were filled in with biotinylated nucleotides, and free blunt ends were ligated. After ligation, crosslinks were reversed and the DNA purified from protein. The DNA was sheared to ~350 bp mean fragment size, and sequencing libraries were generated using NEBNext Ultra enzymes (New England Biolabs, Ipswich, USA) and Illumina-compatible adapters. The libraries were sequenced on an Illumina HiSeqX to produce 310M read pairs. SRA Accession: SRR29099389.

*10x Chromium Linked Reads (2016):* We sent the tissue of a single Alex Yellow female to 10x Genomics for HMW DNA extraction via their "fresh frozen tissue" protocol (https://support.10xgenomics.com/genome-exome/sample-prep/doc/demonstrated-protocol-hmw-dna-extraction-from-fresh-frozen-tissue) followed by their Version 2 Chromium version 2 library preparation and sequencing (https://support.10xgenomics.com/genome-exome/sample-prep/doc/user-guide-chromium-genome-reagent-kit-v2-chemistry). This sequencing produced 347M paired-end reads (SRA Accession: SRR32688197).

*PacBio (2018):* We applied a user-developed protocol for Qiagen genomic-tip DNA extraction for insects (Genomic-tip 100/G) that uses a gravity column for HMW extractions. Before extraction we dissected, froze, and then pooled whole-body tissue from the 20 largest then-available adult females from our Alex Yellow line. We added RNase A and Proteinase K to the pooled tissue to degrade non-DNA molecules. Next, we washed the degraded tissue with Buffer QC and QF during column runs in order to separate DNA from cell detritus and eluted the DNA into nanopure water. After extraction, we re-suspended samples in 120 µL of T-low-E buffer (10mM Tris pH 7.9, 0.1mM EDTA) prior to quality checks and library preparation. We then used a NanoDrop 1000 to assess the quality of the re-suspended sample; our extraction garnered ~100 µg of DNA with a 260/280 of 1.90 and a 260/230 of 2.36.

We then sent the HMW DNA to the Arizona Genomics Institute for sequencing. Pacbio sequencing library construction followed the Pacbio 20-kb protocol. The ligated molecules were size-selected at 10 kb with Blue Pippin. The resulting library was bound with sequencing primer v3 and sequencing polymerase v2, followed by loading to 1M Sequel SMRT cells in the Pacbio Sequel instrument. Nine cells were sequenced from library concentration loadings between 15 pmol to 20 pmol. CLR reads are available at SRA (SRR29047464), as are the CCS reads (SRR29047459).

*Transcriptome sequencing:* We used bulk RNA-Seq of various tissues and whole individuals from the Alex Yellow sexual inbred line to annotate the *P. antipodarum* genome assembly. All laboratory lineages were maintained in standard conditions (e.g., [5]) in the Neiman lab snail room until dissection. For all samples, dissected tissues to be used for RNA extraction were immediately placed in Qiagen RNA*later*® following manufacturer protocols and then frozen at -80ºC until RNA extraction. RNA was extracted with Trizol and cleaned with Zymo RNA Clean and Concentrate following manufacturer protocols. We produced cDNA libraries for sequencing with the TruSeq RNA Sample Prep v2. All transcriptomes were sequenced with Illumina HiSeq 2500 2x100 bp sequencing.

*Data Preprocessing:*
*Read processing and quality control:* All Illumina data were processed using the FASTQ Quality Trimmer (FASTX-Toolkit) [6] with the Illumina parameter set to 33 and a quality threshold of 20. We included all reads of zero length. We then used the FastQC to ensure that the quality of the reads was increased after trimming. PacBio continuous long read (CLR) subreads were extracted from HDF5 movie files using bax2bam software (Pacific Biosciences, San Diego, USA).

#### Assembly and annotation of the *P. antipodarum* genome

*Genome size estimation with flow cytometry:* We determined approximate total nuclear genome size using flow cytometry, as described in [7]. Briefly, flash-frozen *P. antipodarum* and *P. estuarinus* head tissues were ground in a solution containing 0.2 M Tris-HCl (pH 7.5), 4 mM $MgCl_2$, 1% TritonX-100, and 4 µg/mL DAPI. This solution was filtered through a 70-micron nylon sheet and assayed on a Beckman-Coulter Quanta SC MPL flow cytometer (Beckman Coulter, Brea, USA). Four *P. estuarinus* samples were assayed on a run on January 27th, 2015, and 40 *P. antipodarum* Alex Yellow samples were assayed on February 13th, 2015. We used the FL1 channel to assess DAPI fluorescence (and thus the total DNA content) of cell nuclei under a UV lamp. At the beginning and end of each flow cytometry run, we calibrated the machine using 20 µL of chicken red blood cells (Lampire Biological Labs, Pipersville, PA) treated and filtered as for *Potamopyrgus* samples. Each sample was run until a count of at least 10,000 events was achieved.

*Contig assembly and scaffolding:* Our first-pass assembly of PacBio CLR reads was carried out using the Canu assembler v.1.7.1 [8]. We used two different sets of parameters for this assembly step. First, we generated a ‘collapsed’ or ‘haploid’ assembly in which the assembly was collapsed into a single haplotype by relying upon default assembly parameters associated with the ‘pacbio’ CLR parameter set. Second, we attempted to avoid collapsing paralogous regions of the genome to produce an ‘uncollapsed’ or ‘polyploid’ assembly using the following flags: *corOutCoverage=200 "batOptions=-dg 3 -db 3 -dr 1 -ca 500 -cp 50"*. While we had relatively high-quality sequencing of the focal genome, we polished ‘collapsed’ and ‘uncollapsed’ contigs separately using the consensus caller Arrow (Pacific Biosciences, San Diego, USA) for PacBio CLR data first, then Pilon [9] for Illumina data second, both with default parameters. Following polishing, Dovetail scaffolded ‘collapsed’ contigs using their proprietary HiRise scaffolder (Cantata Bio, Scotts Valley, USA). ‘Uncollapsed’ contigs were not scaffolded, so as to retain structural variants that differed between subgenomes. We then closed gaps and iteratively scaffolded the ‘collapsed’ scaffolds by mapping the scaffolds to themselves to identify overlaps with minimap2 v2.24-r1122 [10], then calling consensus sequences and gap filling with Racon v1.5.0 [11]. Once assembly contiguity asymptoted (six rounds of Racon), we scaffolded the collapsed scaffolds using the 10x Genomics Chromium reads in ARCS v1.05 [12] with default parameters, except for the following: -e 80000, -D enabled, -s 96, -c 4, -l 5, and -B 90 and LINKS v1.8.6, with the following non-default parameters: -a 0.5, and -k 20. We polished scaffolds with Pilon as before, but using the 10x Genomics Chromium linked reads this time, then filled gaps with the PacBio CLR reads using GapFiller v1.10 (BaseClear, Leiden, Netherlands), with the -d flag set to 100, the -i flag set to 10, and all other parameters kept as default. Next, we scaffolded using the small number of circular consensus sequences (CCS) available from the PacBio CLR reads using SSPACE-LongRead v1.1 (BaseClear, Leiden, Netherlands) [13], with default parameters. We repeated the iterative Racon scaffolding and gap-filling, the 10x scaffolding with ARCS and LINKS, polishing with Pilon, gap-filling with GapFiller (CLR reads), and scaffolding with SSPACE-LongRead (CCS reads) one more time. We then finished the assembly with one final round of Racon gap-filling (assembly overlaps) and Pilon polishing (HiSeq Illumina reads). Assembly completeness was evaluated both in terms of sequence and biological content.

*Individual transcriptome generation and analysis:* In order to produce a gene expression set across sexes and female reproductive status, we isolated one brooding adult female (embryos in brood sac, SRA Accession: SRR29099382), one non-brooding adult female (no embryos, SRA Accession: SRR29099381), and one adult male from the Alex Yellow sexual inbred line (SRA Accession: SRR29099380). Once these individuals were identified, we dissected away and stored the body tissue (*i.e.*, containing gonads) in RNA*later*® and at -80ºC until library preparation. We then used these three RNA extractions for library preparation. We barcoded the libraries and ran the three samples split across two separate lanes of an Illumina HiSeq 2000.

*Transcriptome assembly:* We separately assembled transcriptomes for the brooding female, non-brooding female, and male individual *P. antipodarum* with Trinity v2.5.5 [14], used TransDecoder (part of Trinity pipeline) to extract open reading frames, and then CD-HIT-EST v4.6.8 [15] to cluster sequences. Finally, these three transcriptomes were concatenated along with the previously assembled pooled Alex Yellow transcriptome from brooding female ovaries [16]], then we reduced redundancy again with CD-HIT-EST. We included these pooled RNA-sequencing data to enhance and refine annotation of genes related to reproductive function.

*Masking of transposable elements*: We used several *de novo* approaches to identify and assemble transposable elements (TEs) to mask the *P. antipodarum* genome for genome annotation. First, we used RepeatModeler2 v2.0 [17] with RECON and RepeatScout and the LTR Pipeline Extensions enabled (-LTRStruct) on the collapsed genome assembly. We then ran the resulting consensus TE sequences through the REPET PASTECLASSIFIER pipeline v2.0

[18] for further classification based on similarity with multiple TE databases (Pfam27.0_GypsyDB.hmm, epbase20.05_ntSeq_cleaned_TE.fa, and repbase20.05_aaSeq_cleaned_TE.fa). We used SINE_Scan v1.1.1 [19] for *de novo* discovery of SINEs, which are often difficult to identify because of their small size and lack of coding or structural features. We collected the PASTEC classified sequences and SINEs and used them as verified input (--curatedlib) in EDTA v1.8.5 [20]. Finally, we used RepeatMasker [21] to mask the genome assembly with the combined TE library output from EDTA. We used this masked assembly as input for MAKER.

*Gene prediction:* Following the *P. estuarinus* and *P. kaitunuparaoa* genome assembly and annotation [22], we used MAKER v2.31.9 [23] to predict genes in both the collapsed and uncollapsed assembly versions of the *P. antipodarum* genome. Briefly, we ran MAKER iteratively three times with AUGUSTUS v2.5.5 [24] trained before the second and third MAKER runs with BUSCO v3.0.2 (--long --augustus_parameters='--progress=true') based on the transcripts from the previous MAKER output. The first round of MAKER used evidence from multiple *P. antipodarum* transcriptomes assemblies and protein sequences from *Pomacea canaliculata* (GCF_003073045.1 [25]) and *Oncomelania hupensis* (coding sequence from assembled transcriptome SRR1284718, see [22]), and *P. estuarinus* protein sequences [22]. The second and third rounds of MAKER used SNAP v0.15.4 [26] and AUGUSTUS v3.3 to generate the final gene models.

*Functional annotation:* To add functional information to *P. antipodarum* gene predictions, we used the MAKER protein predictions from the un-collapsed assembly to search for protein matches and GO terms in multiple databases. We used DIAMOND BLASTP v2.0.15 (--strand both --query-cover 0.5 --evalue 1e-5) [27] to search for matches in the NCBI nr protein database (downloaded locally, last modified: 2024-02-07 10:05). Additionally, we used DIAMOND BLASTP to query against UniProtKB (release 2024_02) TrEMBL and Swiss-Prot databases. Finally, we used interproscan 5.68-100 [28] to search for GO terms (Date of run: 11/28/2024).

*Meiosis gene inventory:* We inventoried a set of meiosis genes to identify the presence or absence of genes that are critical for sex in *P. antipodarum* and compare the inventory with the meiosis gene inventory from congeners of *P. antipodarum* [22] and with *Pom. canaliculata* (GCF_003073045.1 [25]) and *Marisa cornuarietis* (GCA_004794655.1 [25]). While the genome assemblies presented are from sexual individuals, these data provide a benchmark for direct comparisons of the rate and pattern of evolution of meiosis genes between the many and separate transitions of sexual to asexual reproduction in *P. antipodarum* lineages [29].

*Assembly completeness:* To evaluate the biological completeness of both collapsed and uncollapsed genome assemblies, we used BUSCO v5.4.2 [30] and the metazoa_odb10 lineage reference gene set. We then used the BlobToolKit pipeline [31] with default parameters to summarize assembly statistics and identify potential contamination within the assemblies, visualized in snail plots and blob plots. The BlobTools datasets shown were created using results from BUSCO, BWA-aligned sequencing reads [32], NCBI's blastn (BLAST v2.11.0+) alignments, and DIAMOND blastx v2.0.13 alignments of the genome against the NCBI nucleotide database and the NCBI non-redundant (nr) protein database with taxonomic affiliations, respectively.

*Meiosis gene inventory:* In addition to the meiosis gene inventory methods of [22], we also employed miniprot [33] to determine gene presence using default settings and with the gene models provided by [22] as queries. Miniprot is a splice-aware protein-to-genome aligner that is especially useful for non-model organisms. Of particular relevance is that genes containing exons that are so small that they would not attain the threshold bitscore for discovery in BLAST are able to be located and annotated *via* miniprot. However, because miniprot penalizes non/mis-sense mutations, we still needed to rely on BLAST to extract those genes with signatures of pseudogenization.

For this inventory, we were especially interested in finding genes that exist in multiple copies in the uncollapsed genome assembly because these duplicates are an expected outcome of WGD. We first separated our focal genes in two groups: genes that are used solely during meiosis (*i.e.*, sex-specific genes), and genes that are pleiotropic with respect to functions that are non-meiotic (*i.e.*, genes used in meiosis and mitosis). Next, for genes to be considered present, we set a benchmark of >80% match to the length of the query so that genes that are highly fragmented/incomplete are considered absent. For each gene that is present in at least two copies and has intact reading frames (*i.e*,. miniprot-generated gene models), we aligned the copies using MUSCLE [34] and then used MEGA to calculate the p-distance of nucleotides between the copies [35].

Because a duplicated region in a genome assembly can be an artifact of misassembly in low-complexity regions, for each duplicated gene modeled from miniprot, we confirmed that the apparent mutational differences of each gene copy were reflected in the Illumina reads. We aligned reads to the uncollapsed genome assembly using BWA v0.7.17-r1188 [32] and visually confirmed that the Illumina reads used to generate the assembly matched the unique mutations in each duplicated gene model using the Integrated Genome Viewer [36].

*Site-Frequency Spectrum Analysis:* We compared the SFS of diploid *P. estuarinus* and *P. kaitunuparaoa* to the SFS of presumed diploid *P. antipodarum* to describe the distinct allele frequency distributions for each assembly and whether those distributions are consistent within genomes (*i.e.*, each within-assembly scaffold/contig has the same copy number). We used the sam filtered bam files as described above as input to nQuire with the 'create' command with all settings kept as default except increasing the quality score threshold from 10 to 30 (parameter 'c') to estimate accurate allele counts. We filtered the allele counts using the 'denoise' command with default settings. This tool reduces noise using a Gaussian Mixture Model (GMM) by applying a uniform-noise component to scale-down noise generated by improper mapping and make allele frequency peaks identifiable. Finally we ran the 'lrdmodel' command in nQuire to compare the $\Delta \log L$ free model to each of the three fixed models (diploid, triploid, tetraploid). We ran this program on each of the three genome assemblies as a whole (returning a single best-fit model across all scaffolds and contigs) and on each contig/scaffold within a genome assembly to detect chromosome-level differences in copy number.

*Gene-based tests for genome-duplication:* To infer genome-wide orthologs and paralogs in the *P. antipodarum*, *P. kaitunuparaoa*, and *P. estuarinus* genomes, we used OrthoFinder v2.5.5 [37] and the -og flag, which outputs homologous gene groups (i.e., "orthogroups") for all input proteomes. This analysis was carried out for primary-transcript-only proteomes of *P. antipodarum* (uncollapsed), *P. kaitunuparaoa,* and *P. estuarinus.* We determined the composition of each orthogroup and binned orthogroups depending on the number of sequences from each species (e.g., single-copy genes – 1:1:1; genes duplicated in *P. antipodarum*, but single-copy in the other species – 2:1:1; genes that are multi-copy in *P. antipodarum*, but single-copy in other species – N:1:1; *etc.*). The number of orthogroups falling into the bins with species-specific duplicates was compared to test the hypothesis of a historical WGD event in *P. antipodarum*. Specifically, we used a Fisher's Exact test to test whether the *P. antipodarum* genome contained more gene duplicates than would be expected by gene duplication alone. This was accomplished by comparing the number of orthogroups that contained two sequences from *P. antipodarum*, but only a single sequence from each of the other two species (*i.e.*, 2:1:1 orthogroup bin) to the number of analogous orthogroups for the other two species (*i.e.*, 1:2:1 and 1:1:2 orthogroup bins). These orthogroups were also used in downstream synteny- and phylogeny-based analyses.

Whole-genome duplication events result in the simultaneous duplication of every gene in the genome, leading to the expectation that divergence post-WGD should exhibit a unimodal distribution with a peak in divergence around the relative time of the event. The primary alternative scenario that could also explain our previous observations (i.e., increased nuclear genome DNA content, the prevalence of skewed SFS at biallelic sites, and the overrepresentation of orthogroups with gene duplicates of *P. antipodarum* relative to its congeners), would be many individual gene or segmental duplications (or reduction in the rate of gene duplicate purging) in *P. antipodarum*. In this alternative scenario, individual gene duplicates would be expected to have no global temporal signature of divergence between paralogs because each duplication event would have arisen independently from all others. The expected result is that divergence between paralogs should be uniformly distributed, with no distinct peaks (assuming a constant rate of individual gene duplication). In addition, tree topology can be used to infer the mechanism of gene number increase. First, a lineage-specific WGD event (i.e., autopolyploidy) would be expected to result in a sister relationship between homoeologs, to the exclusion of sequences from other species, in a majority of gene trees. An allopolyploidization event in which two separate lineages hybridized to form the third polyploid lineage would be expected to show homoeologs grouping with the diploid that is most closely related to the progenitor species, assuming progenitor lineages had been sampled. In the ‘many individual gene duplication events’ scenario, paralog relationships would depend upon the timing of the gene duplication events, such that no enrichment for a particular topology (i.e., paralogs sister to one another; one paralog grouping with *P. kaitunuparaoa*; or one paralog grouping with *P. estuarinus*) would be expected. We estimated phylogenetic distance and inferred tree topology for all orthogroups featuring a lineage-specific duplicate (i.e., 2:1:1, 1:2:1, and 1:1:2 orthogroup bins).

To estimate the distribution of phylogenetic distance between duplicate genes and compare gene tree topologies, we generated multiple-sequence alignments for all orthogroups, using MAFFT v7.525 [38] to align protein sequences and the align_fasta_with_mafft_codon perl subroutine found in sloan.pm (https://github.com/dbsloan/perl_modules/blob/master/sloan.pm) to back-translate those alignments into CDS sequences, as in [39]. We trimmed CDS alignments using the protein-based trimming algorithm in ClipKIT v2.3.0 [40], using the -l flag to log the trimmed positions. Trimmed amino acid alignments were then back-translated into CDS alignments using a custom Python script designed to interpret the ClipKIT logfile (https://github.com/jsharbrough/protTrim2CDS). Individual gene trees (for orthogroups with more than 3 sequences) were then inferred in RAxML v8.2.12 [41] using the raxmlHPC-PTHREADS implementation of the rapid Bootstrap analysis and search for best-scoring ML tree (‘-f a’ flag), with the model specified as “GTRGAMMAIX” and a random seed generated for each orthogroup separately, using 100 bootstrap replicates. Because RAxML does not work for alignments with fewer than three sequences, we estimated phylogenetic distance for all sequences in orthogroups with three sequences present (*i.e.*, the 1:1:1 orthogroup bin) using PhyML v3.3.20180214 [42], with the following parameter set: ‘-b 100 -m 012345 -f m -v e -c 4 -a e -o tlr -s SPR --n_rand_starts 5 --rand_start --r_seed <random_seed>’. Tree topology was assessed for all trees in the 2:1:1, 1:2:1, and 1:1:2 orthogroup bins using a custom Python script (https://github.com/jsharbrough/potamomics) that leverages the DendroPy package [43]. Briefly, tree topology was first determined using an unrooted approach by checking for one of two possible topologies (*i.e.*, *P. antipodarum* sequences group together or one *P. antipodarum* sequence groups with a *P. estuarinus* or *P. kaitunuparaoa* sequence). Trees were then midpoint-rooted to determine the relative root placement on each tree. The same script also extracted branch lengths from trees to calculate patristic distance between all sequences in the orthogroup. We also assessed midpoint-rooted tree topology and patristic distance in single-copy genes (*i.e.*, 1:1:1 orthogroups) using a custom Python script (https://github.com/jsharbrough/potamomics) to evaluate relationships within the *Potamopyrgus* genus as in [22].

To test whether the duplicated genes in *P. antipodarum* exhibited conserved synteny and to better understand the genome structure, we qualitatively evaluated duplicate collinearity in the *P. antipodarum* uncollapsed genome assembly compared to the assemblies from the other species and to the collapsed assembly. Our expectation was that if a WGD had occurred in *P. antipodarum*, gene duplicates would display collinearity in the uncollapsed assembly, whereas an elevated rate of individual gene duplication would not be expected to generate any such pattern. We manually generated collinearity data from the positional information found in the assembly GFF files and combined these data with orthogroup compositional information described above. We then graphically plotted these data for each contig, using scaffolds from the collapsed assembly as anchors, in R [44]. The R code underlying these synteny plots was generated in collaboration with ChatGPT-4-turbo (openai.com) and is available at (https://github.com/jsharbrough/potamomics).

*Transposable element identification and comparative analyses:* With recent advances in automated curation of transposable elements, we sought to simplify our TE discovery and annotation method for improved continuity and reproducibility. Therefore, we conducted a second round of TE sequence discovery for comparative analyses. The portion of the genome masked by the first and second versions of our TE libraries are nearly identical.
We identified TEs from the un-collapsed *P. antipodarum* genome assembly with RepeatModeler v2.0.5 [17], using rmblast 2.14.1+ as the search engine, TRF 4.09, RECON 1.08, and RepeatScout 1.0.6 for repeat consensus building, and LTR_Retriever v2.9.0. for LTR structural analysis. We then ran the RepeatModeler output consensus sequences through Manual Curation Helper (MCHelper) v1.6.6 for automated TE library curation (e.g., reducing redundancy, extending TE consensus sequences to improve completeness, and incorporating structural features into classification)[45]. We then used RepeatMasker v4.1.7-p1 [21] with NCBI/RMBLAST v. 2.14.1+ as the search engine to characterize the TE content of the *P. antipodarum* uncollapsed and collapsed assemblies and the *P. estuarinus* assembly [22]. Following repeat masking, we used the RepatMasker utility scripts buildSummary.pl and calcDivergenceFromAlign.pl to summarize the results and collect Kimura substitution distances for TE families, which we visualized as repeat landscapes with R [44].

To directly compare the repetitive portion of the *P. antipodarum* genome to that of its congeners *P. estuarinus* and *P. kaitunuparaoa*, we used DNApipeTE v.1.3.1_07.dec.2017 [46] on the reads from the *P. antipodarum* Yellow lineage reads to assemble repeats and estimate their genomic abundance. As in Fields et al. (2024) [22], we used a coverage of 0.2x, along with an approximate 900Mb genome size based on flow cytometric data (Supplementary Figure S1 and [47]) and the uncollapsed genome assembly size. To calculate comparable per-TE family abundance estimates across these three species, we used deviaTE [48], which maps genomic sequence reads to a TE library for depth-based abundance measurements. We used a set of single-copy sequences [49] as internal controls to estimate per-haploid copy numbers. We evaluated TEs in terms of relative abundance between each of the possible pairwise comparisons across the three species by dividing the TE copy number of one species by the other species for every TE family. These relative values helped to differentiate TEs with similar and different abundances between the three species. Additionally, we used RepeatProfiler v1.1 [50] to generate comparative visualizations of TE abundance and intragenomic variation (e.g., substitutions between copies of elements in the same family), similarly based on mapping reads to the TE library and single copy sequence panel.

*Note on transposable element classification:* The MCHelper classification of transposable elements largely follows the hierarchical  scheme outlined in by Wicker et al. (2007) [51] At the highest classification level, TEs are considered either Class I retrotransposons that reverse transcribe an intermediate RNA molecule during replication (i.e., “copy-and-paste”), or Class II DNA transposons that do not – most have TIRs and rely on transposase for excision and reintegration elsewhere (i.e., “cut-and-paste”), but other non-RNA elements, like rolling-circle based helitrons, are also Class I. The structural features and replicative mechanism of retrotransposons are used to further divide Class I – largely – into LTR and non-LTR, the long and short interspersed nuclear elements, LINEs and SINEs, respectively. Another important TE distinction that may or may not factor into its classification is between autonomous elements, which are capable of self-replication, and nonautonomous elements, which rely on an autonomous element to recognize – often through some sequence homology – and replicated it, e.g., SINEs rely on LINEs for replication.

### Results

##### Assembly and annotation of the *P. antipodarum* genome

*Assembly:* Our completed uncollapsed *P. antipodarum* assembly had a contig N50 of 38.1k and total length of 993M (Figure 1B, Supplementary Figure S2A). The collapsed version of our completed *P. antipodarum* assembly had a scaffold N50 of 1.14M and total length of 543M (Supplementary Figure S2B). The ~83% larger uncollapsed assembly appears to better reflect the genome of *P. antipodarum* than the collapsed assembly because it is largely consistent with DAPI-based flow cytometric estimates of total nuclear genome size obtained from *P. antipodarum* (~2.06 pg) and *P. estuarinus* (~1.26 pg) (Figure 1C, Supplementary Figure S1), showing that *P. antipodarum* has a substantially larger nuclear genome than its congeners (also see [47]).

*Annotation:* In total, we annotated 57,703 protein-coding genes in the uncollapsed contigs, which was 79.0% more genes than the *P. estuarinus* assembly (32,237) and 90.4% more genes than the (relatively incomplete) *P. kaitunuparaoa* assembly (30,310) [22]. By contrast, we annotated 30,790 protein-coding genes in the *P. antipodarum* collapsed scaffolds. Our functional annotation analysis returned 40,850 *P. antipodarum* genes from the uncollapsed assembly with matches to the NCBI nr database, 39,015 to the UniProt TrEMBL database, 22,618 to the UniProt Swiss-Prot database, and 22,241 with GO terms from InterPro. We also characterized the functional completeness of both the uncollapsed and collapsed assemblies using BUSCO v.5.4.2 [30] and the metazoa_odb10 reference gene, finding that 90.8% of the 954 reference BUSCOs were complete, 43.7% were complete in multiple copies, 3.9% were fragmented, and 5.3% were missing in the uncollapsed assembly (Figure 1B, Supplementary Figure S2A). In the collapsed assembly, 84.8% of the 954 reference BUSCOs were complete, 4.2% were complete with multiple copies, 6.2% were fragmented, and 9% were missing (Supplementary Figure S2B). Following this initial BUSCO analysis, we also used the Blobtools pipeline [31] to quantify potential contamination in both assemblies (Supplementary Figure S3).

##### Comparative transposable element abundances in *Potamopyrgus*

The highest copy number measured for each species by deviaTE was the same element, a 498 bp sequence classified as an LTR retroelement by MCHelper. The estimated copy numbers for this TE were 2,748 in *P. antipodarum* and 3,892 and 3,404 in *P. estuarinus* and *P. kaitunuparaoa*, respectively. The variant frequency profiles from RepeatProfiler for this TE were nearly identical across in the three *Potamopyrgus* species for this element, indicating that its activity occurred largely or even exclusively before these three species split and may longer be active (Figure S8).

The most extreme disparity in relative TE abundance for *P. antipodarum* vs. *P. estuarinus* (1337.6x higher) and *P. kaitunuparaoa* (1145.2x higher), was a DIRS retroelement that is apparently present only in the *P. antipodarum* genome, at an estimated 17 copies. The thousandfold-higher estimated copy number is likely an outcome of a few reads mapping nonspecifically in *P. estuarinus* and *P. kaitunuparaoa*, leading to estimated copy numbers of 0.014 and 0.015 respectively, which likely reflects absence of this element in their genomes (Figure S9). The RepeatProfiler output for this element reveals that only a handful of substitutions have occurred across the nearly 6 Kb length of the element, indicative of a recent invasion that may have occurred exclusively in *P. antipodarum*.

Based on our RepeatMasker results, the TE with the lowest Kimura substitution level (0.3%) is a terminal-inverted repeat DNA transposon of the Mariner family (based on submission to CENSOR online, https://www.girinst.org/censor/index.php). It is also among the most “complete” elements, indicated by the average length of 1,421 bp for this element compared to the curated RepeatModeler reference length of 1,517 bp. This Mariner element has an open reading frame coding for transposase, indicating it is likely an autonomous element (i.e., capable of self-mobilization). We identified only two nearly full-length copies of this element in both *P. estuarinus* and *P. kaitunuparaoa* genomes but 314 copies in the *P. antipodarum* genome, with nearly all of these copies having identical nucleotide sequences (Supplementary Figure 10). We aligned these 318 sequences with MAFFT v7.505 (automated alignment strategy selection of FFT-NS-i), then trimmed the sequence with trimAl v1.4.rev15 [52] (-automated1), and used IQ-TREE 2.2.2.7 [53] for a maximum likelihood tree reconstruction with 1000 ultrafast bootstraps [54]. The phylogenetic reconstruction of sequences from all three species indicates that *P. antipodarum* shares copies that have higher similarity to sequences from its congeners, and that a single lineage of this transposon expanded to hundreds of copies (Supplementary Figure S10A). The read-alignment visual summarization from RepeatProfiler shows a similar picture, with nearly 100% identity for each position (Supplementary Figure S10B). Most (13/20) of the TEs in *P. antipodarum* with apparent “bursts” of activity – based on relative copy numbers across species – also demonstrated evidence of recent transposon activity based on the RepeatMasker results, reflected in Kimura substitution values below 3%.

Based on our RepeatMasker results, the TE with the longest average genomic copy (8,024 bp) is one classified as a Maverick (also known as Polinton), a TE characterized by its large size and virus-like coding sequences, including polymerase B and integrase – both of which are present in this element [55]. This Maverick has a reference sequence length of 13,140 bp and a Kimura substitution level of 0.5% (third-lowest overall), indicating that many of the copies in the genome are complete and identical, a strong indication of recent activity. Our deviaTE estimates of copy number were 165 for *P. antipodarum*, 3 for *P. estuarinus*, and 2 for *P. kaitunuparaoa*. We obtained 169 full-length copies of this Maverick element from the P. antipodarum genome assembly, 2 from the *P. estuarinus* assembly and none from *P. kaitunuparaoa*.

#### SUPPLEMENTARY TABLES

**Supplementary Table S1.** TE families discovered in *P. antipodarum* uncollapsed assembly using RepeatModeler, with automated curation and classification by MCHelper.

**Supplementary Table S2.** Scaffold-specific *in silico* ploidy estimates from nQuire for 100 longest scaffolds in *P. antipodarum* collapsed genome assembly.

**Supplementary Table S3.** Scaffold-specific *in silico* ploidy estimates from nQuire for 100 longest scaffolds in *P. estuarinus* genome assembly.

**Supplementary Table S4.** Scaffold-specific *in silico* ploidy estimates from nQuire for 100 longest scaffolds in *P. kaitunuparaoa* genome assembly.

**Supplementary Table S5.** Tree topology distributions of gene trees from various orthogroup bins.

**Supplementary Table S6.** Per-TE results for P. antipodarum RepeatMasker results and comparative copy number statistics for 3 *Potamopyrgus* species with DeviaTE

#### SUPPLEMENTARY FIGURE LEGENDS

**Supplementary Figure S1.** Density plots describing distribution of raw DAPI fluorescence intensities from two runs of flow cytometry for *P. estuarinus* (green curves, top) and Alex Yellow *P. antipodarum* (blue curves, bottom). Each curve represents a different sample, with black lines representing chicken red blood cell standards, which we run at the beginning and end of each flow run (solid = beginning of run, dashed = end of run). All four *P. estuarinus* samples exhibited similar genome sizes. There was more variation in genome size in Alex Yellow individuals, including at least two triploid contaminant individuals. The *P. estuarinus* run took place on January 27th, 2015 and the Alex Yellow run took place on February 13th, 2015.

**Supplementary Figure S2.** Snail plots comparing the “uncollapsed” contig assembly (A) and “collapsed” scaffold assembly (B) of *P. antipodarum*. The snail plot in the center of each panel is divided into 1,000 size-ordered bins, with each bin representing 0.1% of the assembly. In both panels, the sequence length distribution is shown in dark gray, with the radius scaled to the longest sequence (shown in red). The N50 sequence length (orange) and N90 sequence length (pale orange) are represented by the arcs in the center of each plot. The cumulative sequence count of each assembly is represented by the pale gray spiral, with white scale lines showing orders of magnitude. GC and AT percentages are reflected by the dark blue and pale blue areas, respectively, around the outside of the plot. N percentages are reflected in white areas in the “collapsed” scaffold assembly. A summary of BUSCO markers from the metazoa_odb10 set found to be complete, fragmented, duplicated, and missing in each assembly is shown in the top right of each panel.

**Supplementary Figure S3.** Comparison of sequence coverage and length between “uncollapsed” and “collapsed” assemblies of *P. antipodarum*. The blob plots show the base coverage against GC proportion for sequences in the “uncollapsed” contig assembly (A) and “collapsed” scaffold assembly (B), with histograms showing the sequence length sum distribution along each axis. Circles are sized in proportion to sequence length on a logarithmic scale. Plots C and D show the cumulative sequence length for the “uncollapsed” contig assembly (C) and the “collapsed” scaffold assembly (D). The cumulative lengths of all sequences assigned to each phylum via NCBI taxonomic hits are indicated by colored lines. In all plots, sequences are colored by phylum.

**Supplementary Figure S4**. nQuire output from the 100 scaffolds from the *P. antipodarum* collapsed assembly with the largest number of biallelic sites. One scaffold did not favor either model.

**Supplementary Figure S5.** Upset plot describing orthologous gene group composition in *P. antipodarum*, *P. kaitunuparaoa*, and *P. estuarinus*. *Top* – Histogram of orthogroup number within bins. The blue column represents the 2:1:1 orthogroup bin, for which *P. antipodarum* has two gene copies, while *P. kaitunuparaoa* and *P. estuarinus* have only a single copy. Green columns represent analogous orthogroups in which *P. kaitunuparaoa* and *P. estuarinus* have double copy genes, but the other species only have a single copy. *Bottom* – Orthogroup bins describing gene content in which white circles represent single-copy genes, black circles represent double-copy genes, and black circles encircling white circles represent multi-copy (i.e., >2) genes. Absence of a circle indicates the absence of a gene from that particular species.

**Supplementary Figure S6.** Densitree depicting branch lengths and tree topologies of gene trees in the 2:1:1 orthogroup bin, whose *P. antipodarum* sequences were sister to one another.

Each individual gene tree is represented by a thin-lined tree, with the median tree branches being represented in the background by the thick solid lines. For the sake of visual clarity, only those trees whose max root-to-tip distance was ≤ 0.1 are shown here (n = 1693/2113), but all 2113 trees were included in median tree calculation. The four observed tree topologies are depicted in each panel. The top left panel represents the primary hypothesis for the species tree. The top right panel represents the current hypothesis for the introgression tree (i.e., introgression between *P. estuarinus* and *P. kaitunuparaoa,* see Fields et al., 2024). The bottom left represents the other ILS tree, and the bottom right represents either ancient paralogs or introgression with rate heterogeneity among *P. antipodarum* homoeologs.

**Supplementary Figure S7.** Global patterns of microsynteny across the *P. antipodarum* genome. Each panel represents an individual scaffold from the collapsed assembly that contained ≥1 orthogroup from the 2:1:1 orthogroup bin. Scaffolds (middle row) were used as anchor points for evaluating microsynteny in uncollapsed contigs (top and bottom rows). Black lines connect genes (colored bars) from the same orthogroup. Gene rearrangements in one subgenome vs. the other are visible in the form of black lines that cross other black lines. Most scaffolds exhibited few if any rearrangements of these double-copy genes, which are still maintained in duplicate in the *P. antipodarum* genome. Rearrangements are more prevalent in the single-copy genes (Figure 5), suggesting that recombination has played a prominent role in gene loss and rediploidization in *P. antipodarum*.

**Supplementary Figure S8.** RepeatProfiler visualization output for high copy transposable element (TE) in *Potamopyrgus*. Top three rows show read depth, normalized by single copy sequences, across the element, color coded according to normalized depth per-base. Bottom three rows show un-normalized read depth per-base, with color coded substitutions relative to reference TE sequence. This TE (rnd-1_family-444_unconfirmed) is classified as a Class I LTR retrotransposon by MCHelper. It has the highest estimated copy number for all three *Potamopyrgus* species.

**Supplementary Figure S9.** RepeatProfiler visualization output for DIRS retrotransposon found exclusively in *P. antipodarum*. Top three rows show read depth, normalized by single copy sequences, across the element, color coded according to normalized depth per-base. Bottom three rows show un-normalized read depth per-base, with color coded substitutions relative to reference TE sequence. This TE (rnd-5_family-4408) is classified as a Class I DIRS (family *DIRS*) retrotransposon by MCHelper. “This graph has no coverage” annotation for *P. estuarinus* and *P. kaitunuparaoa* due to low coverage.

**Supplementary Figure S10.** Burst of *Mariner* DNA transposon activity in *P. antipodarum*. (A) Maximum-likelihood molecular phylogenetic tree (IQ-TREE2, substitution model = K2P; log-likelihood = -9313.030) of approximately full length copies from uncollapsed *P. antipodarum* genome assembly (314 sequences), *P. estuarinus* assembly (2 sequences) and *P. kaitunuparaoa* assembly (2 sequences). (B) RepeatProfiler visualization output, with sequence schematic depicting transposase open reading from above. Top three rows show read depth, normalized by single copy sequences, across the element, color coded according to normalized depth per-base. Bottom three rows show un-normalized read depth per-base, with color coded substitutions relative to reference TE sequence. This TE (rnd-5_family-12008_inc) is classified as a Class II TIR DNA transposon by MCHelper.

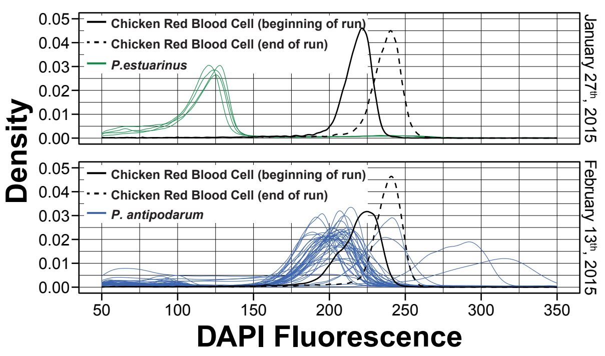

Density
DAPI Fluorescence
Chicken Red Blood Cell (beginning of run)
Chicken Red Blood Cell (end of run)
P.estuarinus
January 27th, 2015
Chicken Red Blood Cell (beginning of run)
Chicken Red Blood Cell (end of run)
P. antipodarum
February 13th, 2015
0.00
0.01
0.02
0.03
0.04
0.05
50
100
150
200
250
300
350

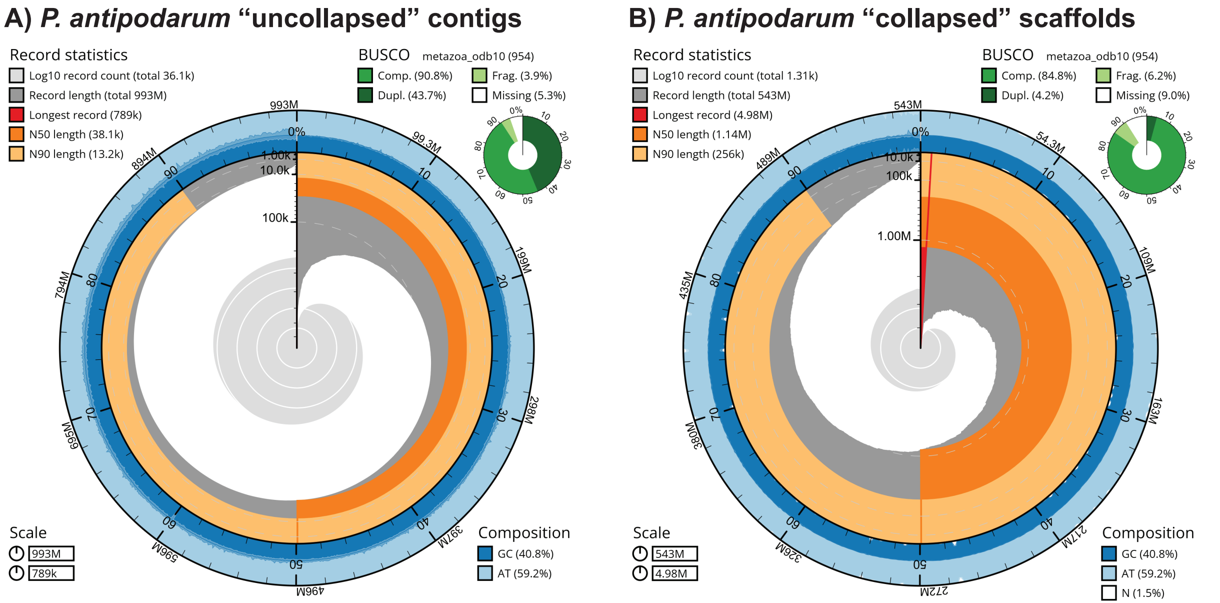
A) P. antipodarum "uncollapsed" contigs
Record statistics
Log10 record count (total 36.1k)
Record length (total 993M)
Longest record (789k)
N50 length (38.1k)
N90 length (13.2k)
BUSCO metazoa_odb10 (954)
Comp. (90.8%)
Frag. (3.9%)
Dupl. (43.7%)
Missing (5.3%)
Scale
993M
789k
Composition
GC (40.8%)
AT (59.2%)
B) P. antipodarum "collapsed" scaffolds
Record statistics
Log10 record count (total 1.31k)
Record length (total 543M)
Longest record (4.98M)
N50 length (1.14M)
N90 length (256k)
BUSCO metazoa_odb10 (954)
Comp. (84.8%)
Frag. (6.2%)
Dupl. (4.2%)
Missing (9.0%)
Scale
543M
4.98M
Composition
GC (40.8%)
AT (59.2%)
N (1.5%)

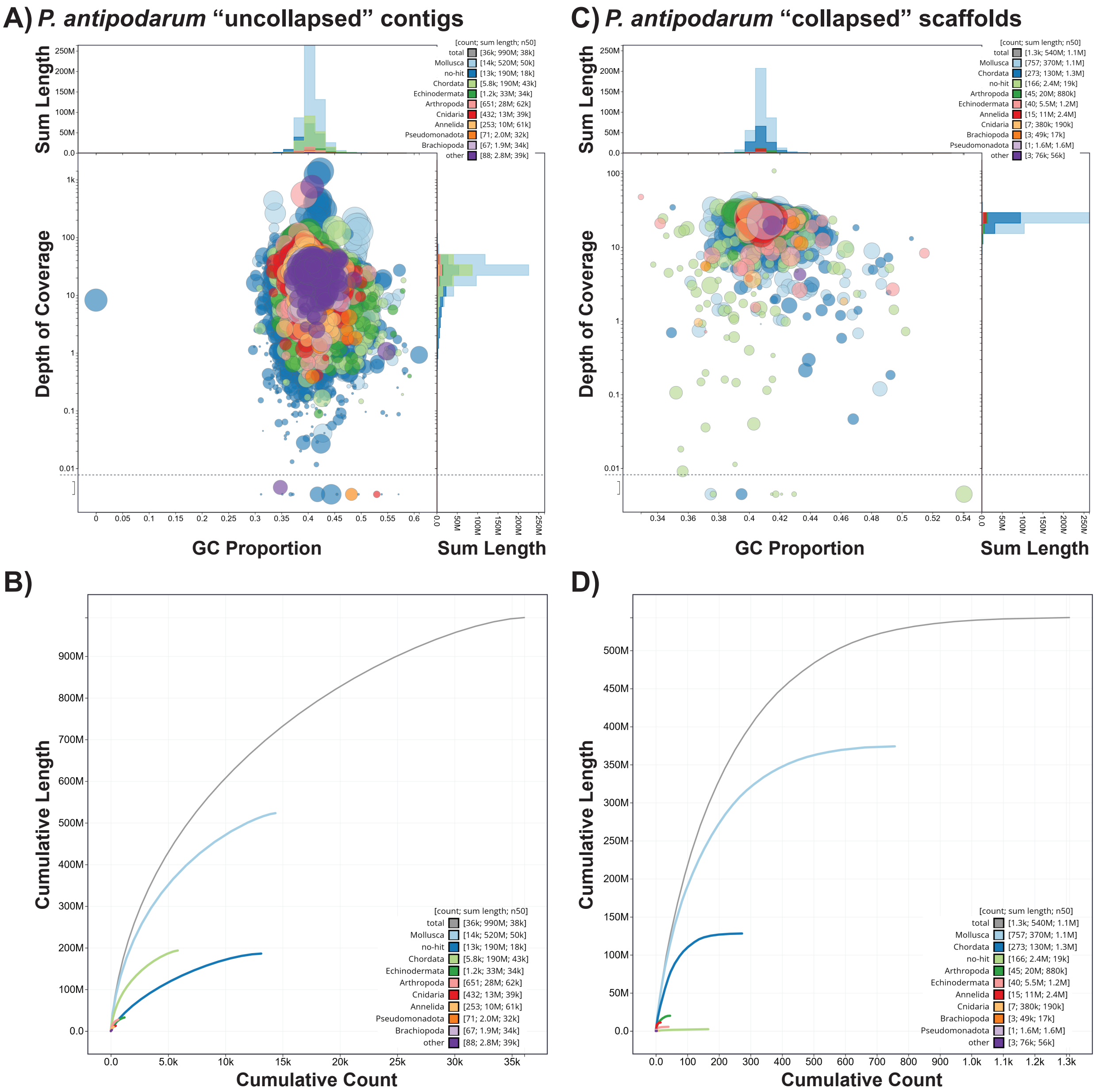
A) P. antipodarum "uncollapsed" contigs
C) P. antipodarum "collapsed" scaffolds
Sum Length
Depth of Coverage
GC Proportion
[count; sum length; n50]
total [36k; 990M; 38k]
Mollusca [14k; 520M; 50k]
no-hit [13k; 190M; 18k]
Chordata [5.8k; 190M; 43k]
Echinodermata [1.2k; 33M; 34k]
Arthropoda [651; 28M; 62k]
Cnidaria [432; 13M; 39k]
Annelida [253; 10M; 61k]
Pseudomonadota [71; 2.0M; 32k]
Brachiopoda [67; 1.9M; 34k]
other [88; 2.8M; 39k]
[count; sum length; n50]
total [1.3k; 540M; 1.1M]
Mollusca [757; 370M; 1.1M]
Chordata [273; 130M; 1.3M]
no-hit [166; 2.4M; 19k]
Arthropoda [45; 20M; 880k]
Echinodermata [40; 5.5M; 1.2M]
Annelida [15; 11M; 2.4M]
Cnidaria [7; 380k; 190k]
Brachiopoda [3; 49k; 17k]
Pseudomonadota [1; 1.6M; 1.6M]
other [3; 76k; 56k]
B)
D)
Cumulative Length
Cumulative Count

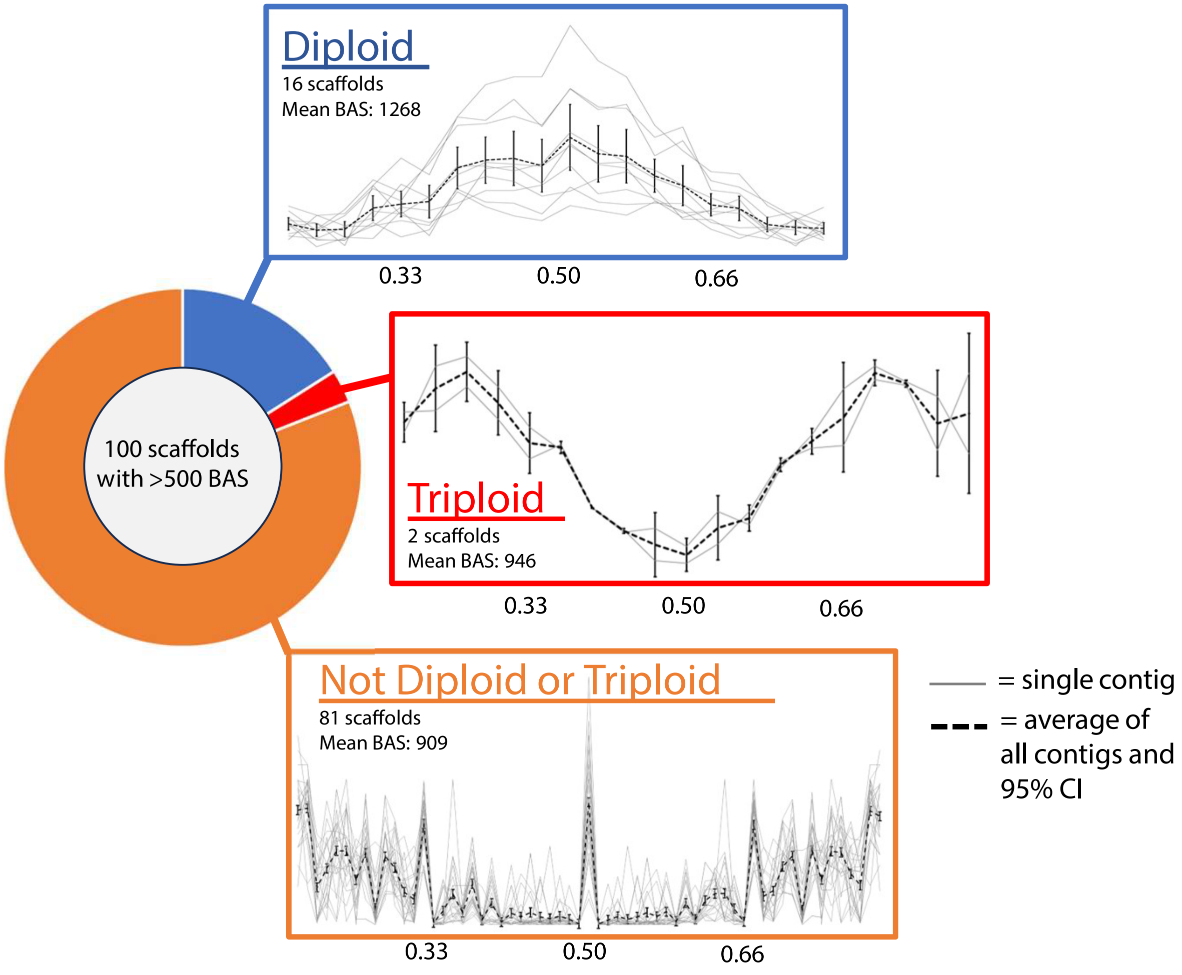
Diploid
16 scaffolds
Mean BAS: 1268
0.33
0.50
0.66
100 scaffolds
with >500 BAS
Triploid
2 scaffolds
Mean BAS: 946
0.33
0.50
0.66
Not Diploid or Triploid
81 scaffolds
Mean BAS: 909
0.33
0.50
0.66
= single contig
= average of all contigs and 95% CI

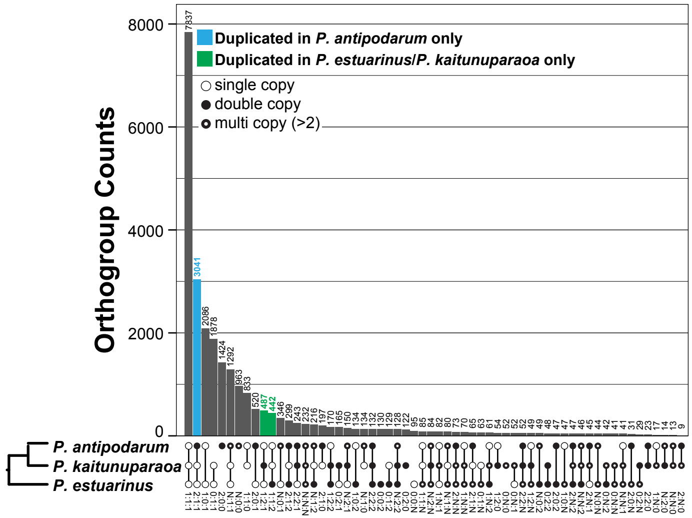

Orthogroup Counts
8000
6000
4000
2000
0
Duplicated in *P. antipodarum* only
Duplicated in *P. estuarinus*/*P. kaitunuparaoa* only
single copy
double copy
multi copy (>2)
7837
3041
2086
1878
1424
1292
963
833
520
487
442
346
299
243
232
216
197
170
165
150
134
134
132
130
129
128
122
95
85
84
82
80
73
70
65
63
61
54
52
52
52
49
49
48
47
47
47
46
45
44
42
41
41
31
29
23
17
14
13
9
*P. antipodarum*
*P. kaitunuparaoa*
*P. estuarinus*

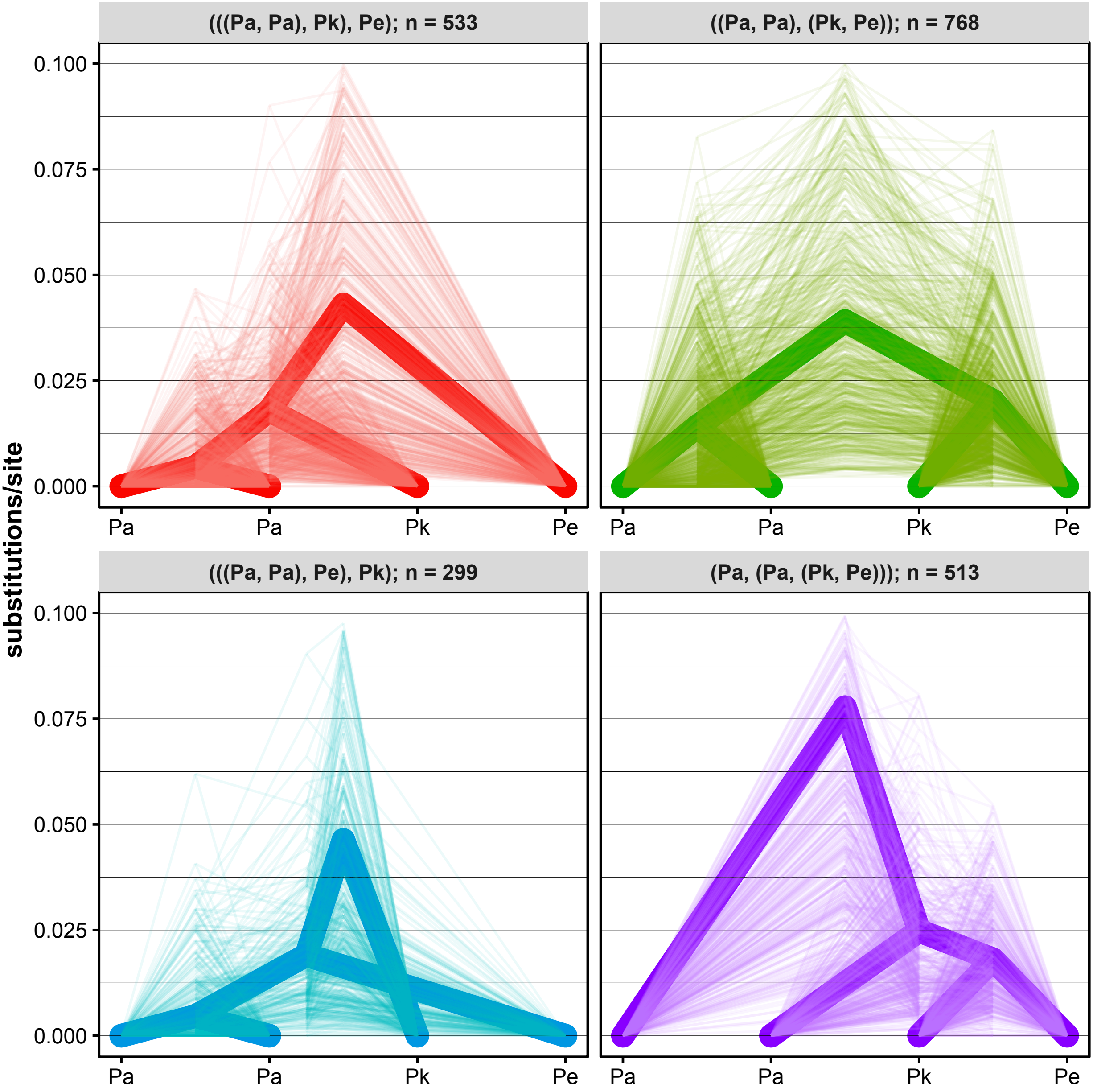

(((Pa, Pa), Pk), Pe); n = 533
((Pa, Pa), (Pk, Pe)); n = 768
(((Pa, Pa), Pe), Pk); n = 299
(Pa, (Pa, (Pk, Pe))); n = 513
substitutions/site
0.100
0.075
0.050
0.025
0.000
Pa
Pa
Pk
Pe

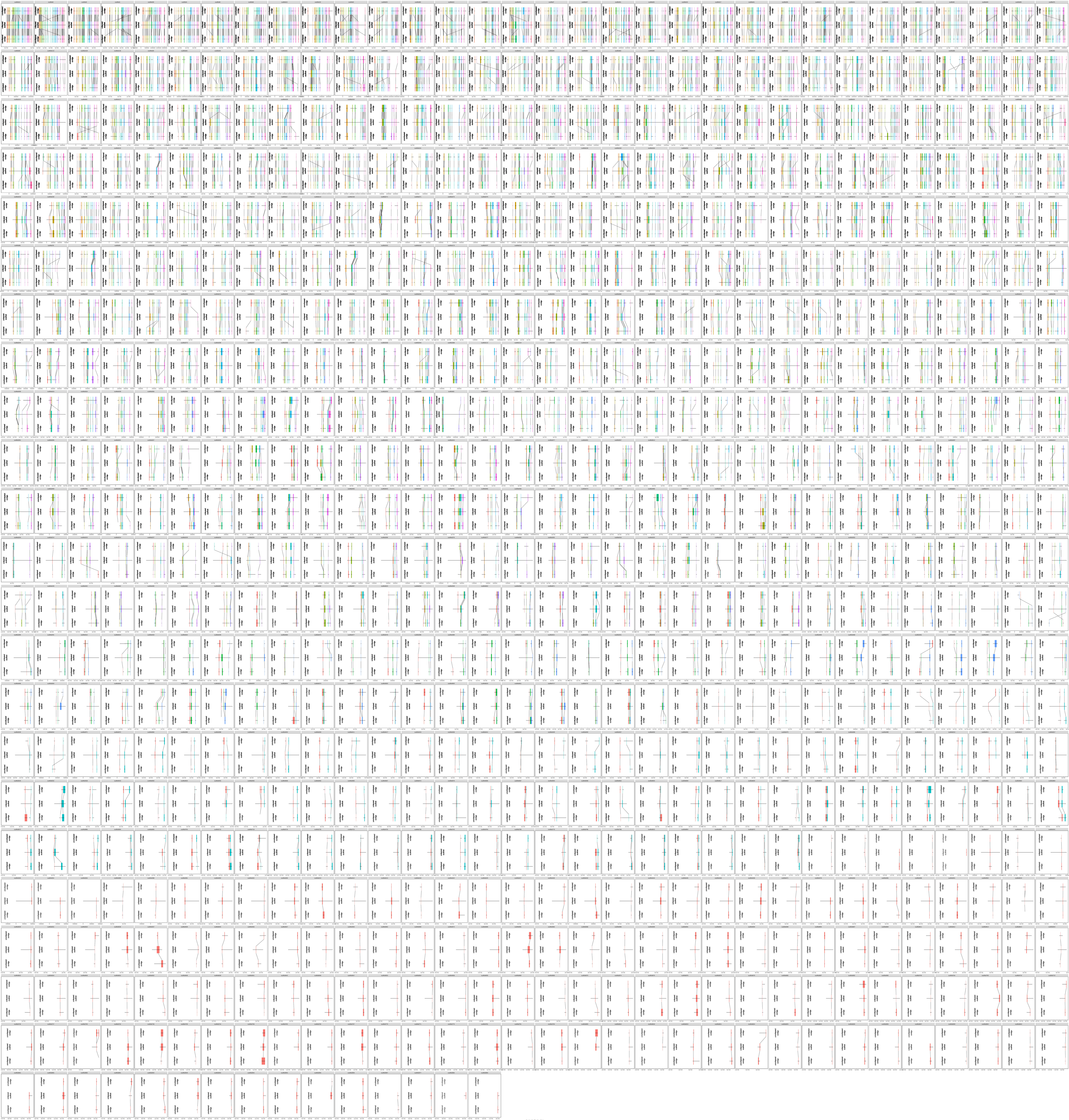

Nucleotide Position (bp)

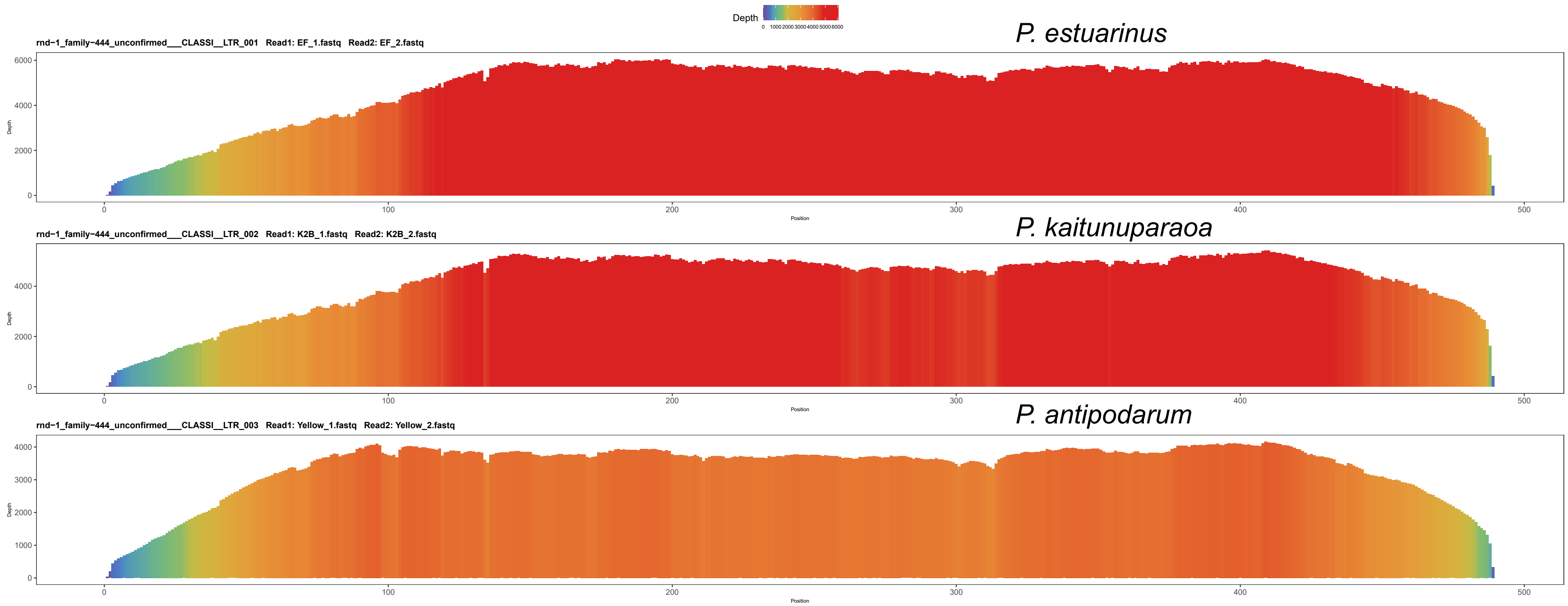
Depth
0 1000 2000 3000 4000 5000 6000
P. estuarinus
rnd−1_family−444_unconfirmed___CLASSI__LTR_001 Read1: EF_1.fastq Read2: EF_2.fastq
P. kaitunuparaoa
rnd−1_family−444_unconfirmed___CLASSI__LTR_002 Read1: K2B_1.fastq Read2: K2B_2.fastq
P. antipodarum
rnd−1_family−444_unconfirmed___CLASSI__LTR_003 Read1: Yellow_1.fastq Read2: Yellow_2.fastq
Depth
Position

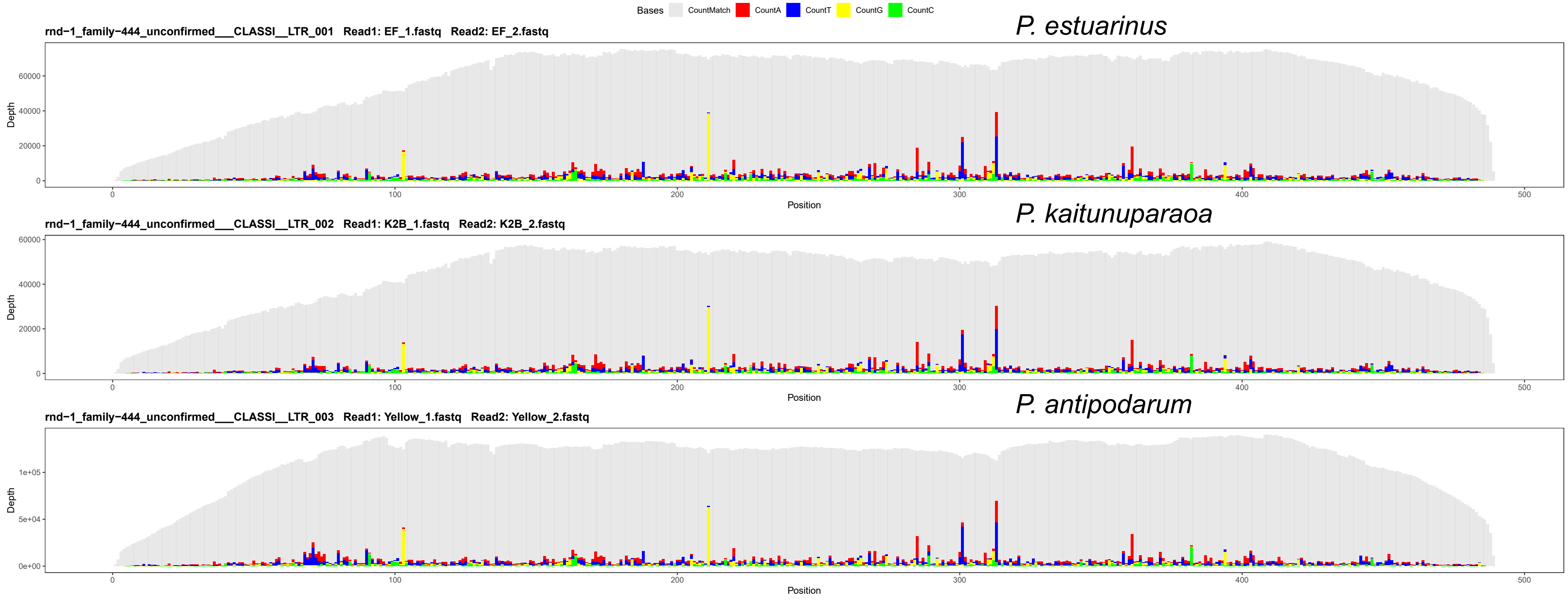
Bases CountMatch CountA CountT CountG CountC
P. estuarinus
rnd−1_family−444_unconfirmed___CLASSI__LTR_001 Read1: EF_1.fastq Read2: EF_2.fastq
P. kaitunuparaoa
rnd−1_family−444_unconfirmed___CLASSI__LTR_002 Read1: K2B_1.fastq Read2: K2B_2.fastq
P. antipodarum
rnd−1_family−444_unconfirmed___CLASSI__LTR_003 Read1: Yellow_1.fastq Read2: Yellow_2.fastq
Depth
Position

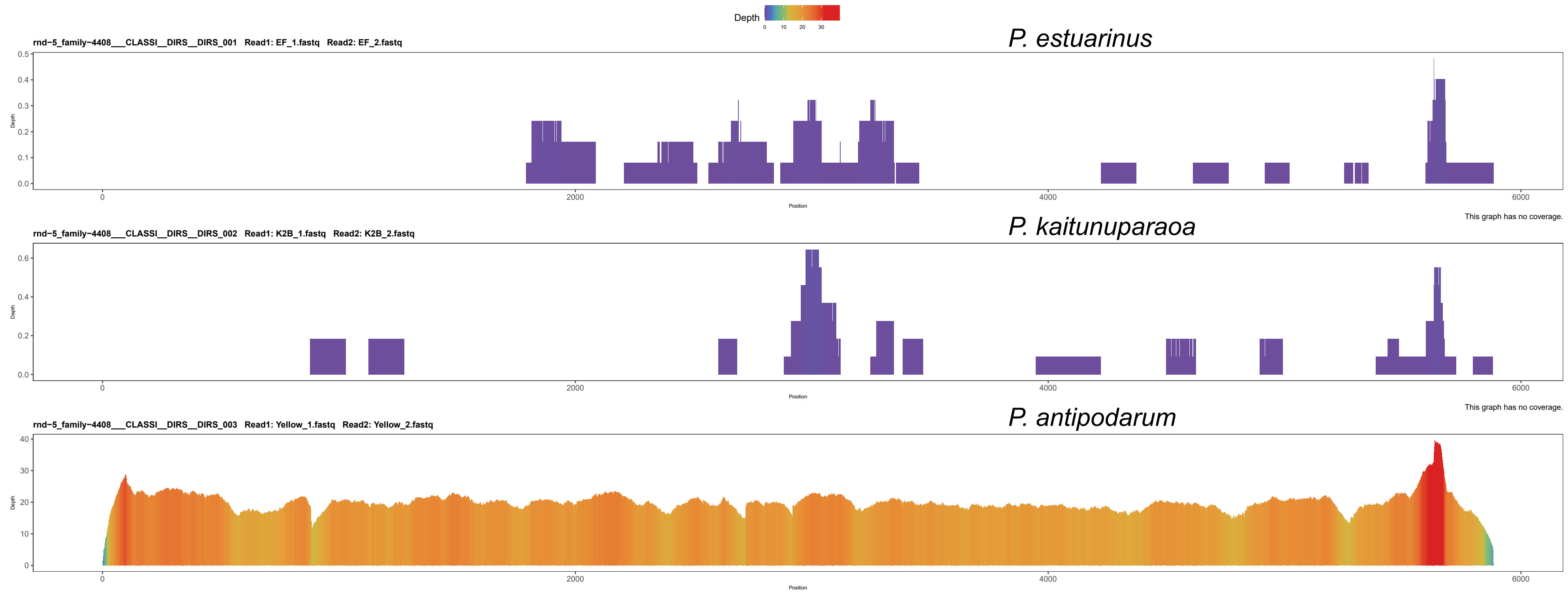
Depth
0
10
20
30
P. estuarinus
rnd-5_family-4408___CLASSI__DIRS__DIRS_001 Read1: EF_1.fastq Read2: EF_2.fastq
This graph has no coverage.
P. kaitunuparaoa
rnd-5_family-4408___CLASSI__DIRS__DIRS_002 Read1: K2B_1.fastq Read2: K2B_2.fastq
This graph has no coverage.
P. antipodarum
rnd-5_family-4408___CLASSI__DIRS__DIRS_003 Read1: Yellow_1.fastq Read2: Yellow_2.fastq
Depth
Position
0
2000
4000
6000

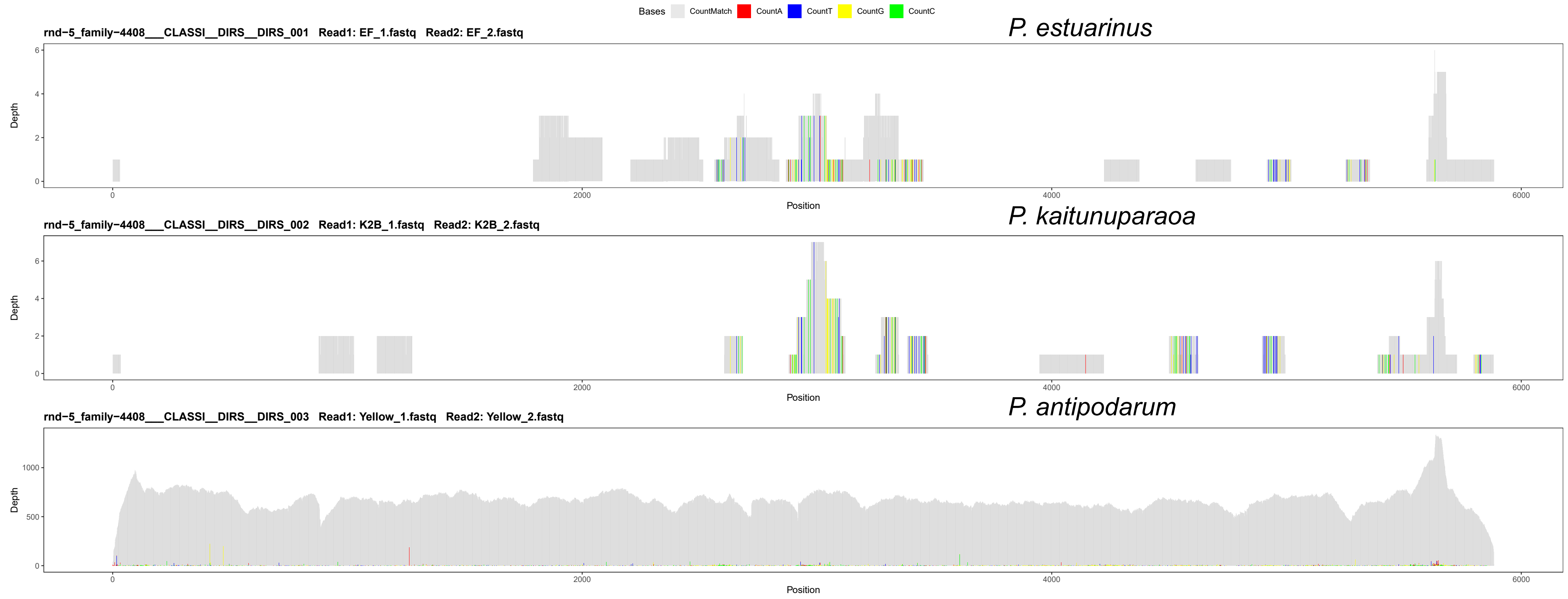
Bases
CountMatch
CountA
CountT
CountG
CountC
P. estuarinus
rnd-5_family-4408___CLASSI__DIRS__DIRS_001 Read1: EF_1.fastq Read2: EF_2.fastq
P. kaitunuparaoa
rnd-5_family-4408___CLASSI__DIRS__DIRS_002 Read1: K2B_1.fastq Read2: K2B_2.fastq
P. antipodarum
rnd-5_family-4408___CLASSI__DIRS__DIRS_003 Read1: Yellow_1.fastq Read2: Yellow_2.fastq
Depth
Position
0
2000
4000
6000

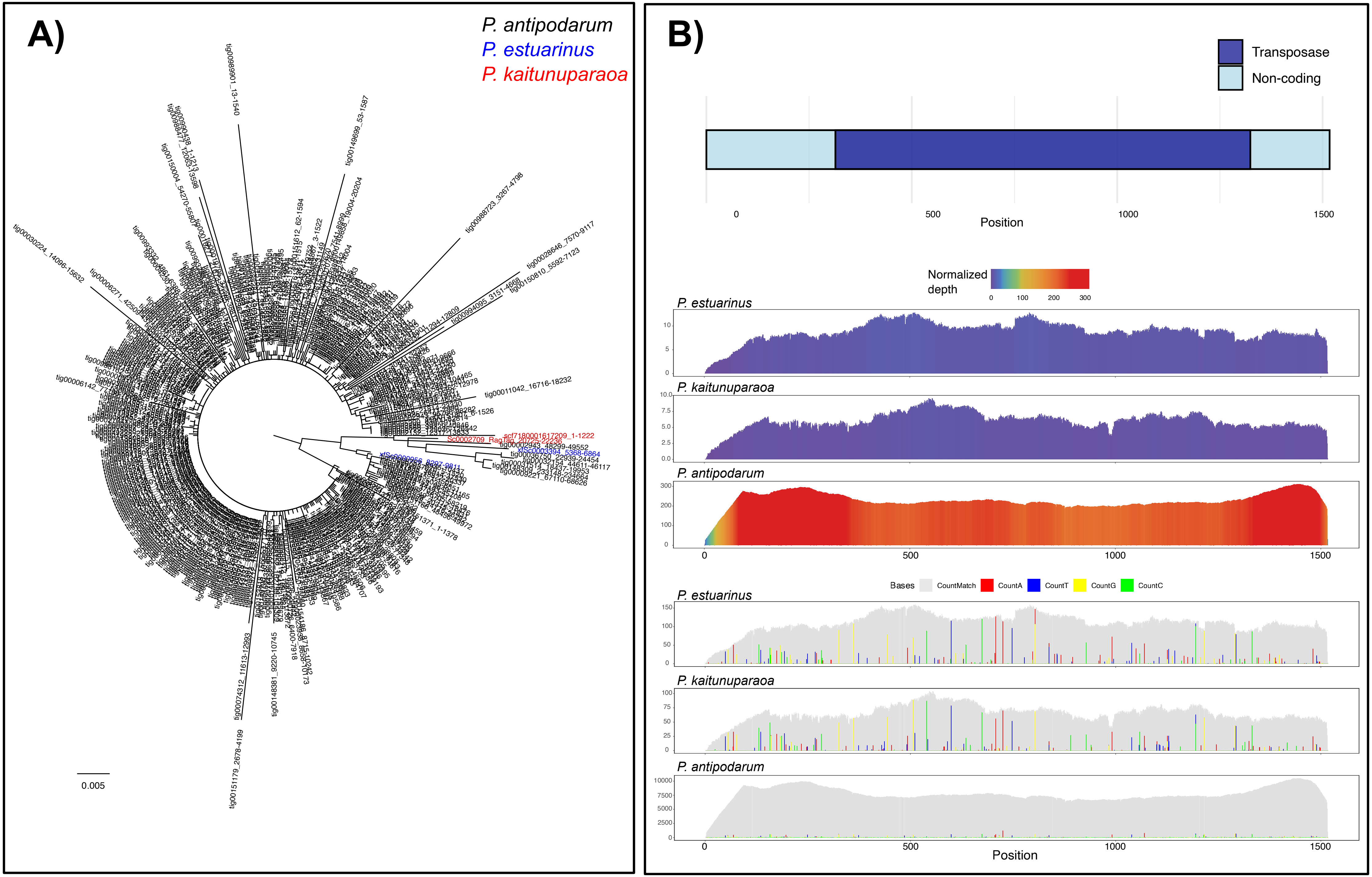
A)
P. antipodarum
P. estuarinus
P. kaitunuparaoa
0.005
B)
Transposase
Non-coding
0
500
1000
1500
Position
Normalized depth
0
100
200
300
P. estuarinus
P. kaitunuparaoa
P. antipodarum
Bases
CountMatch
CountA
CountT
CountG
CountC
P. estuarinus
P. kaitunuparaoa
P. antipodarum
Position
